# Discovery of novel ID2 antagonists from pharmacophore-based virtual screening as potential therapeutics for glioma

**DOI:** 10.1101/2021.05.10.443505

**Authors:** Genshen Zhong, Yichun Wang, Qi Wang, Minna Wu, Yichuang Liu, Shitao Sun, Zhenli Li, Jinle Hao, Peiyuan Dou, Bin Lin

**Affiliations:** Henan Key Laboratory of Immunology and Targeted Therapy, Henan Collaborative Innovation Center of Molecular Diagnosis and Laboratory Medicine, School of Laboratory Medicine, Xinxiang Medical University, Xinxiang, Henan, 453003, China; School of Basic Medicine, Xinxiang Medical University, Xinxiang, Henan, 453003, China; School of Pharmaceutical Engineering, Shenyang Pharmaceutical University, Shenyang, Liaoning, 110016, China; School of Chemistry, Cardiff University, Cardiff, CF10 3AT, United Kingdom

**Keywords:** glioma, inhibitor of differentiation, pharmacophore, virtual screening

## Abstract

Glioma, especially the most aggressive type glioblastoma multiforme, is one of the central nervous system malignant cancer with a poor prognosis. Traditional treatments are mainly surgery combined with radiotherapy and chemotherapy, which is still not satisfactory. Therefore, it is of great clinical significance to find new therapeutic agents. Served as an inhibitor of differentiation, protein ID2 (inhibitor of DNA binding 2) plays an important role in neurogenesis, neovascularization and malignant development of gliomas. It has been shown that ID2 affects the malignant progression of gliomas through different mechanisms. In this study, a pharmacophore-based virtual screening was carried out and 16 hit compounds were purchased for pharmacological evaluations on their ID2 inhibitory activities. Based on the cytotoxicity of these small-molecule compounds, two compounds were shown to effectively inhibit the viability of glioma cells in the low micromolar range. Among them, AK-778-XXMU was chosen for further study due to its better solubility in water. A SPR assay proved the high affinity between AK-778-XXMU and ID2 protein with the KD value as 129 nM. The plausible binding mode in the biding site of ID2 was studied by molecular docking. Subsequently, the cancer-suppressing potency of the compound was characterized both *in vitro* and *in vivo*. The data demonstrated that compound AK-778-XXMU is a potent ID2 antagonist which has the potential to be developed as a therapeutic agent against glioma.

**Highlights:**

- Two pharmacophores were built from the first-in-class pan-ID antagonists AGX51
- A pharmacophore-based virtual screening was carried out and 16 hit compounds were purchased for pharmacological evaluations in glioma inhibition
- Compound AK-778-XXMU was identified to be a potent ID2 antagonist in the low submicromolar range (KD: 159 nM)

## Introduction

Malignant glioma is the most aggressive brain tumor with a high rate of fatality and poor prognosis [1–3]. Despite the enormous efforts invested to improve the efficacy of classical combination of surgical resection supplemented by chemotherapeutic agents, the overall survival of the patients is still not promising [2]. The chemotherapeutic agents currently approved by the FDA for the treatment of malignant glioma include temozolomide (Temodar, TMZ) and bevacizumab (Avastin). Unfortunately, as the first-line drug for glioblastoma, which is the most aggressive type of glioma, the emergence of TMZ resistance has greatly limited the effectiveness of the drug. Bevacizumab is currently the only targeted drug approved by FDA for the treatment of recurrent glioblastoma [4]. It has also been reported that bevacizumab can prolong the progression-free survival of newly diagnosed glioblastoma patients, but the overall survival time cannot be prolonged [5]. Moreover, the discovery of a subtype of tumor cell with stem cell-like self-renewal properties [6], along with insufficient tumor resection due to invasion of normal brain tissue by tumor cells [7] has been shown to result in relatively ineffective treatment of glioma [8]. Therefore, it is of great urgence to develop novel potential therapeutics against novel targets that play key roles in the progression of glioma.

Inhibitors of DNA binding (ID) proteins, commonly expressed in mammals synergistically, regulate gene expression, cell lineage localization and cell differentiation in tissues [9,10]. They belong to the family of helix-loop-helix (HLH) transcription factors. However, the absence of a basic motif which is necessary for binding to DNA distinguishes it from the basic helix-loop-helix transcription factors (bHLH), which contributes to its role of transcription factor inhibitor. Ever since the first ID gene, ID1, was identified in 1990 by Benezra *et al* [9], three more ID groups were discovered and named ID2, ID3, and ID4 [11–14]. The crucial role of the ID proteins is due to their dominant-negative effect when they form heterodimers with other DNA-binding members of the HLH family, therefore disrupting the protein-DNA interaction. Not surprisingly, strong association of ID proteins with tumor angiogenesis has been suggested [15,16]. In particular, the emerging role of ID2 in glioma has attracted much attention [17]. It was found that ID2 was expressed to varying degrees in different grades of glioma and was significantly increased in mesenchymal astrocytoma and glioblastoma compared to low-grade malignant astrocytoma, which was not detected in normal cerebral white matter [18]. Furthermore, the expression level of ID2 was significantly correlated with the survival of glioblastoma cells in a low glucose metabolic stress environment by influencing mitochondrial function [19,20]. During the development of myeloid cells, Kinase insert domain receptor (KDR, a type IV receptor tyrosine kinase), also known as vascular endothelial growth factor receptor 2 (VEGFR-2), a critical gene for VEGF signaling, has the ability to drive differentiation of hematopoietic progenitor cells (HPCs) into tumor-promoting myeloid-derived suppressor cells (MDSCs) which could be enhanced by ID2 by upregulating KDR and then stimulates the formation of an aggressive pro-angiogenic phenotype in gliomas [17,21]. It is believed that neural precursor cells (NPCs), a type of neural stem cells, may also be one origin for glioma cell [22,23]. Phosphorylation of ID2 N-terminal seems to affect its protein expression level in NPCs, in the case of this phosphorylated mutant, ID2 appears to be protected from degradation by the ubiquitin-proteasome system (UPS) to perform a range of pro-oncogenic functions [24].

The ID genes/proteins are critically important during embryonic development but are only active in adults when cancer is present [25–27]. It was found that they are expressed in virtually all cancers [25,28]. The overexpression of ID genes in tumors is linked to an aggressive phenotype and poor clinical outcome. It was proposed that targeting ID proteins was an effective strategy to develop anti-cancer drugs [17].

So far, a variety of strategies have been proposed to interfere with ID activity, including the use of targeted antisense delivery, cell-permeable peptides, antagonists of Id expression, and inhibitors of stability-inducing deubiquitinases, antagonists of ID proteins [29–34]. Among them, the first and only small-molecule pan-ID antagonist is AGX51, discovered by Benezra group [35]. It was shown that AGX51 was able to inhibit Id1-E47 interaction, leading to ubiquitin-mediated degradation of IDs, cell growth arrest, and reduced viability. AGX51 was also able to phenocopy the genetic loss of Id expression in age-related macular degeneration (AMD) and retinopathy of prematurity (ROP) models by inhibiting retinal neovascularization [35]. Due to the therapeutical potential of ID antagonists, it is important to develop more such compounds to target ID proteins.

Therefore, we launched a drug discovery project by utilizing a pharmacophore-based virtual screening approach. Docking-based virtual screening was not chosen because even though ID2 protein has been already solved by crystallography (PDB: 4AYA), the binding site is located at the loop region. Such a region is very flexible in nature so that the binding site may not have a well-defined shape [36]. In contrast, pharmacophore-based virtual screening does not require the structure of the protein [37–40]. The pharmacophore was extracted from AGX51 and subjected to a series of small commercial databases for virtual screening.

In the end, a variety of potential ID2 antagonists were obtained and 16 hit compounds were purchased for pharmacological evaluations for their ID2 inhibitory activities. Based on the cytotoxicity of these small-molecule compounds, two compounds were shown to effectively inhibit the viability of glioma cells at low micromolar concentrations. Among them, AK-778-XXMU was chosen for further study due to its better solubility in water. A SPR (surface plasma resonance) assay proved the high affinity between AK-778-XXMU and ID2 protein with the KD value as 129 nM. Subsequently, the cancer-suppressing potency of the compound was characterized both *in vitro* and *in vivo*. The data demonstrated that compound AK-778-XXMU is a potent ID2 antagonist which has the potential to be developed as a therapeutic agent against glioma.

## Results and Discussion

### 1. Results from pharmacophore-based virtual screening

The commercial compound databases used for virtual screening were CHEMBRIDGE, SPECS, CHEMDIV, ENAMINE, IBS, LIFECHEMICALS, VITAS-M, kindly provided in their electronic form by Topscience Co., Ldt (https://www.tsbiochem.com/). They contained 2,281 compounds in total that were commercially available. Two pharmacophore models (*R*-model and *S*-model) were built from the two enantiomers of AGX51 and they were subjected for virtual screening against those databases mentioned above by using LigandScout 4.0 [38,40]. The screening yielded 24 compounds from the *R*-model and 33 compounds from the *S*-model. After consultation with synthetic chemists for the feasibility of future chemical modifications of those compounds, 16 compounds were purchased. Their chemical structures are listed in **Fig. 1** with their compound names from the respective databases.

**Fig. 1.**
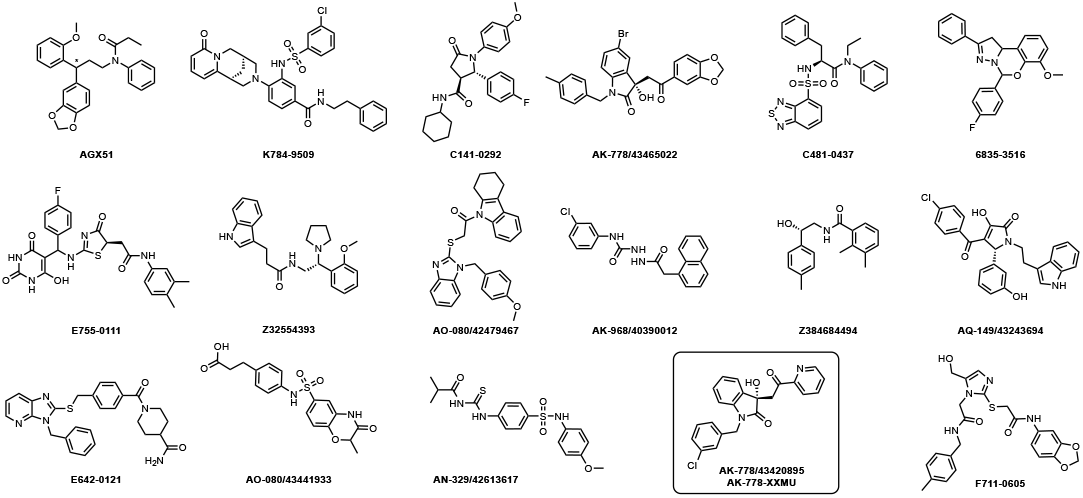
Chemical structures of AGX51 and the 16 hit compounds purchased. The chiral center was labeled with an asterisk. AK-778/43420895 was highlighted in the round rectangle and renamed as AK-778-XXMU for further study.

### 2. Determination of the cytotoxicity of the 16 hit compounds by CCK8 assay

CCK8 assays were conducted next to measure the viability of different glioma cell lines, namely U87, HS683, and GL261, after treated with different concentrations of the 16 hit compounds *in vitro* to examine the cytotoxicity of these compounds against glioma. As shown in **Table 1**, all compounds except three demonstrated detectable cytotoxicity (IC_50_ < 100 μM) against all three glioma cell lines, proving the validity of our pharmacophore-based virtual screening. Among them, compounds AK-778/43465022 and AK-778/43420895 had the lowest IC_50_ values to glioma cells (IC_50_ < 30 μM), indicating they could inhibit cell viability more effectively. Although compound AK-778/43465022 showed the most potent inhibition in all three glioma cell lines, it had poor water solubility **(Fig. 2A)**. Therefore, compound AK-778/43420895 was chosen for further studies. Normal cell lines, human normal hepatocytes (L-02) and mouse embryo fibroblasts (3T3) were exposed to AK-778/43420895 to rule out the potential toxicity and other possible side effects. It was shown in **Fig. 2C** that AK-778/43420895 had little cytotoxicity on normal cell lines. The compound was re-named as AK-778-XXMU for the subsequent studies.

**Table 1.** The cytotoxicity against various glioma cell lines from CCK8 assay.

| Compound Names | $IC_{50}$ ($\mu M$) | | |
| --- | --- | --- | --- |
|  | U87 | HS683 | GL261 |
| K784-9509 | 56.2±3.1 | 55.3±2.1 | 48.1±0.8 |
| E755-0111 | 46.5±1.3 | 40.8±1.5 | 47.9±2.7 |
| C141-0292 | 77.9±3.1 | 84.5±1.7 | 90.2±1.5 |
| E642-0121 | 60.3±1.0 | 62.3±0.4 | 56.7±1.6 |
| Z32554393 | >100 | >100 | 94.6±1.6 |
| AK-778/43465022 | **15.5±0.95** | **18.7±1.3** | **21.3±0.8** |
| AO-080/42479467 | 88.5±0.7 | 91.4±1.3 | 90.4±3.4 |
| AK-968/40390012 | >100 | >100 | >100 |
| AN-329/42613617 | 56.7±4.8 | 53.2±1.7 | 61.3±0.8 |
| AO-080/43441933 | 47.4±1.0 | 41.5±2.6 | 52.5±4.1 |
| C481-0437 | 66.3±1.5 | 69.8±2.6 | 60.6±3.7 |
| 6835-3516 | >100 | >100 | >100 |
| F711-0605 | 37.8±2.7 | 41.5±2.2 | 50.1±1.5 |
| Z384684494 | 67.3±5.0 | 70.3±0.6 | 74.2±1.1 |
| AK-778/43420895 | **27.3±3.7** | **34.2±2.5** | **38.2±5.4** |
| AQ-149/43243694 | 54.3±1.7 | 60.8±0.9 | 57.3±3.1 |

**Fig. 2.**
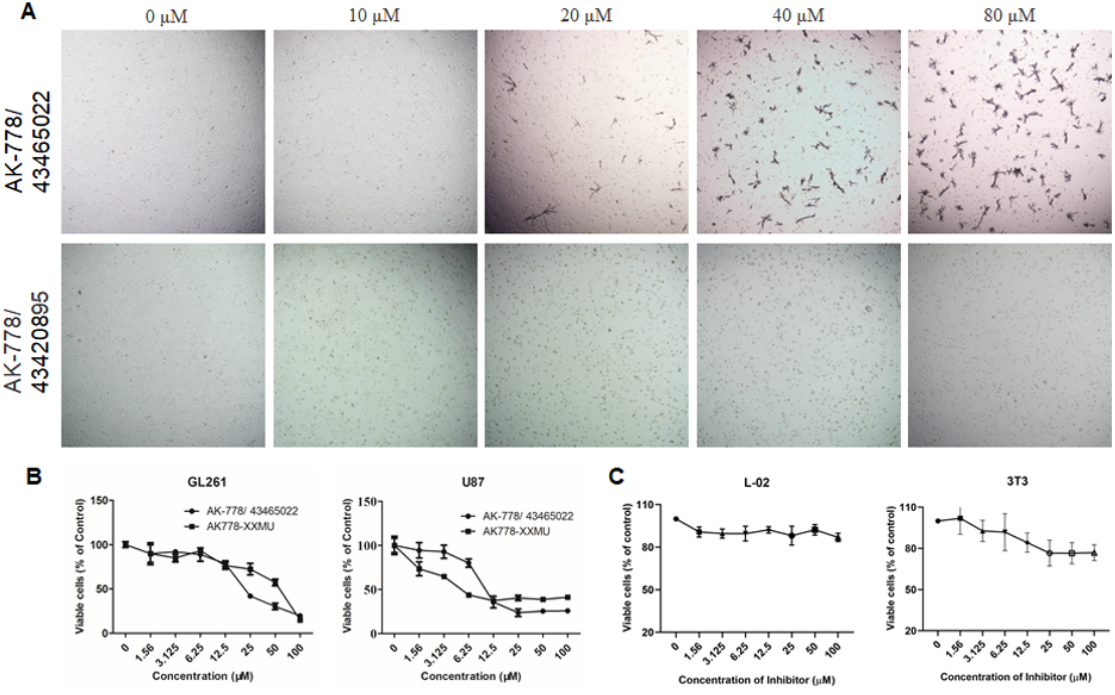
Characterization of compound AK-778/43420895 (AK-778-XXMU) *in vitro*. A: AK-778/43420895 had better water solubility as compared to compound AK-778/43465022. B and C: Compound AK-778/43320895 reduces the viability of glioma cells but not normal cells. The effect of compound AK-778/43320895 on cell viability of Human astroblastoma (U87), mouse glioma (GL261), human normal hepatocytes (L-02) and mouse embryo fibroblasts (3T3) after being exposed to a range of AK-778/43320895 in a concentration gradient for 24 h via CCK8 assay. The percent of cell viability is plotted in a dose-dependent manner. Data points represent percent of cell viability relative to DMSO control of triplicates and expressed as the mean ± SD.

### 3. Determination of the binding affinity between AK-778-XXMU and ID2 protein

Biacore is a new biometric sensing technology based on SPR with advantages of high sensitivity, real-time, fast and label-free, which has been widely used to describe interaction and measure affinity between two partners [41,42]. To identify the quantitative affinity between ID2 protein and compound AK-778-XXMU, SPR assay was performed using a Biacore 8K system (GE Healthcare). ID2 protein was immobilized on the sensor surface as the ligand which was capable of capturing the target compound AK-778-XXMU in solution. By following the kinetics of a binding process of ID2 protein and AK-778-XXMU in real time, the association/disassociation rate constants (ka and kd) and equilibrium disassociation constants (KD) of interactions were obtained. To our delight, the results showed the KD between compound AK-778-XXMU and ID2 protein was 0.129 μM (**Fig. 3**). The submicromolar KD demonstrated that compound AK-778-XXMU was able to bind to ID2 protein with high affinity to exert its inhibitory effects.

**Fig. 3.**
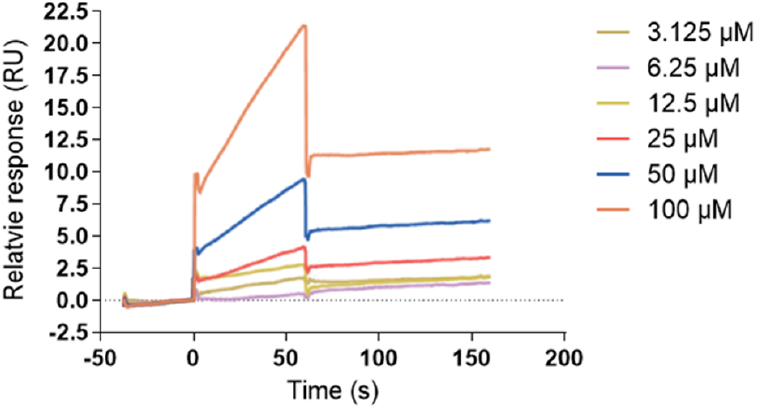
Change in response as a function of time in SPR and the fitting curves of affinity between compound AK-778-XXMU and ID2 at various concentrations. Using 1:1 binding model, ka was determined to be 2.48E+01 (1/Ms), kd was determined to be 3.19E-06 (1/s), and KD was determined to be 1.29E-07 (M). Rmax was 83.6 (RU). Chi^2^ was 0.23 (RU^2^).

### 4. Binding modes of AGX51 and AK-778-XXMU with ID2

Compound AK-778-XXMU was identified from the pharmacophore-based virtual screening described above. The validity of the screening was firmly established by the SPR assay that determined the potent binding of AK-778-XXMU to ID2 with high affinity in the low submicromolar range. The molecular superimposition of AGX51 and AK-778-XXMU as well as their pharmacophores were shown to be nearly perfect, as they should be (**Fig. 4A**). Molecular docking studies were performed to provide more insight into the binding modes of both compounds in the ID2 binding site (**Fig. 4B**). It was shown that both AGX51 and AK-778-XXMU were fit quite well into the hydrophobic crevice in the loop region of ID2 (PDB: 4AYA) [36]. The binding modes of both AGX51 and AK-778-XXMU with ID2 were similar to that of AGX51 with ID1 [35]. As shown in **Fig. 4C** and **Fig. 4D**, in addition to the apparent hydrophobic interactions, a hydrogen bond was observed between the benzodioxole moiety of AGX51 and Gln55. The corresponding hydrogen bond was between the hydroxyl group of AK-778-XXMU and Gln55. Two more hydrogen bonds for AK-778-XXMU were formed between the ketone group and Lys47 as well as between the amide group and Lys58. Pi-cation interactions for both compounds were observed between the two phenyl groups in their structures and Lys47/Lys58. Such interactions offered possible explanations at the atomic level that why AK-778-XXMU was a potent ID2 antagonist.

**Fig. 4.**
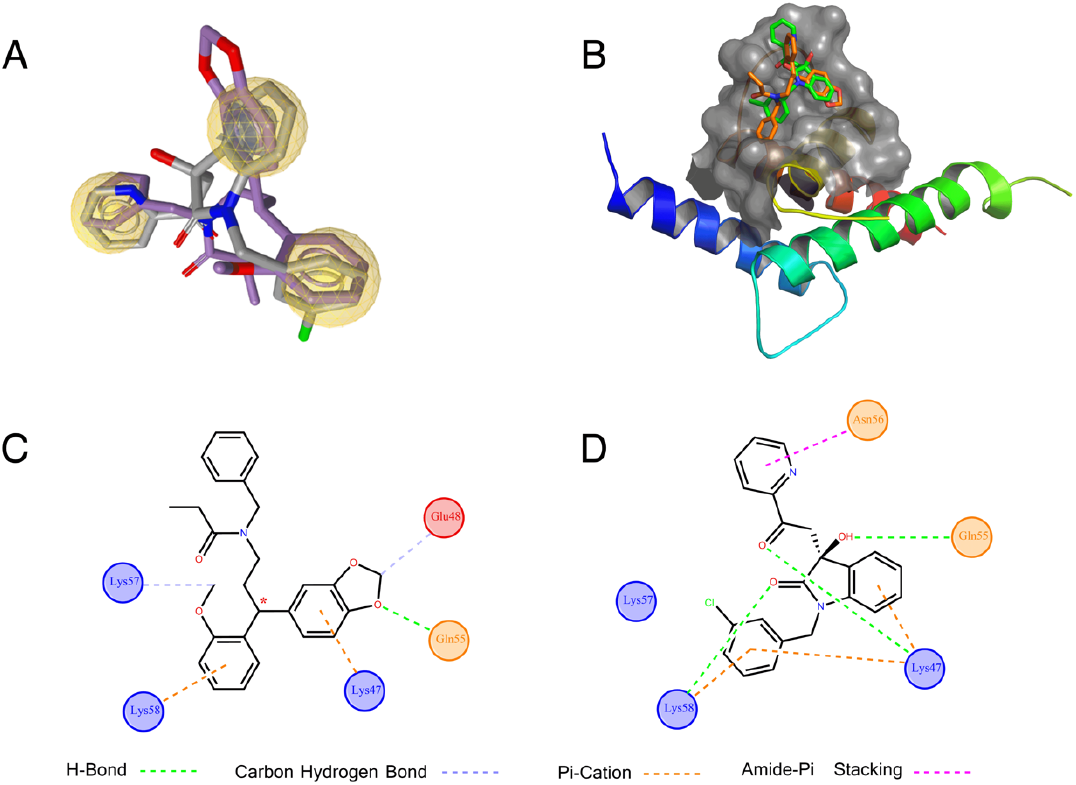
Comparison between AGX51 and the hit compound AK-778-XXMU. (A) molecular superimposition between the S enantiomer of AGX51 (purple) and AK-778-XXMU (grey). Yellow spheres represent hydrophobic centers. (B) AGX51 (orange) and AK-778-XXMU (green) in the binding site of ID2 (C) the 2D diagram of interactions between AGX51 and the binding site of ID2. (D) the 2D diagram of interactions between AK-778-XXMU and the binding site of ID2.

### 5. Effect of compound AK-778-XXMU on cell migration and invasion

To determine whether compound AK-778-XXMU can influence the migration and invasion of glioma cells, wound healing assay was carried out. The results are shown in **Fig. 5**. As expected, compound AK-778-XXMU could slow down glioma cells wound healing. The wound closure rates were depressed significantly as low as (13.11±1.52)% for GL261 and (5.59±1.04)% for U87 after exposed to AK-778-XXMU at the concentration of 12.5 µM. By comparison, closure rates in control group were up to (63.13±0.78)% for GL261 and (61.50±10.21)% for U87 cells. To further verify the effect, the Transwell assay was carried out. As shown in **Fig. 6**, compound AK-778-XXMU could inhibit the migration and invasion of GL261 and U87 cells in a concentration-dependent manner.

**Fig. 5.**
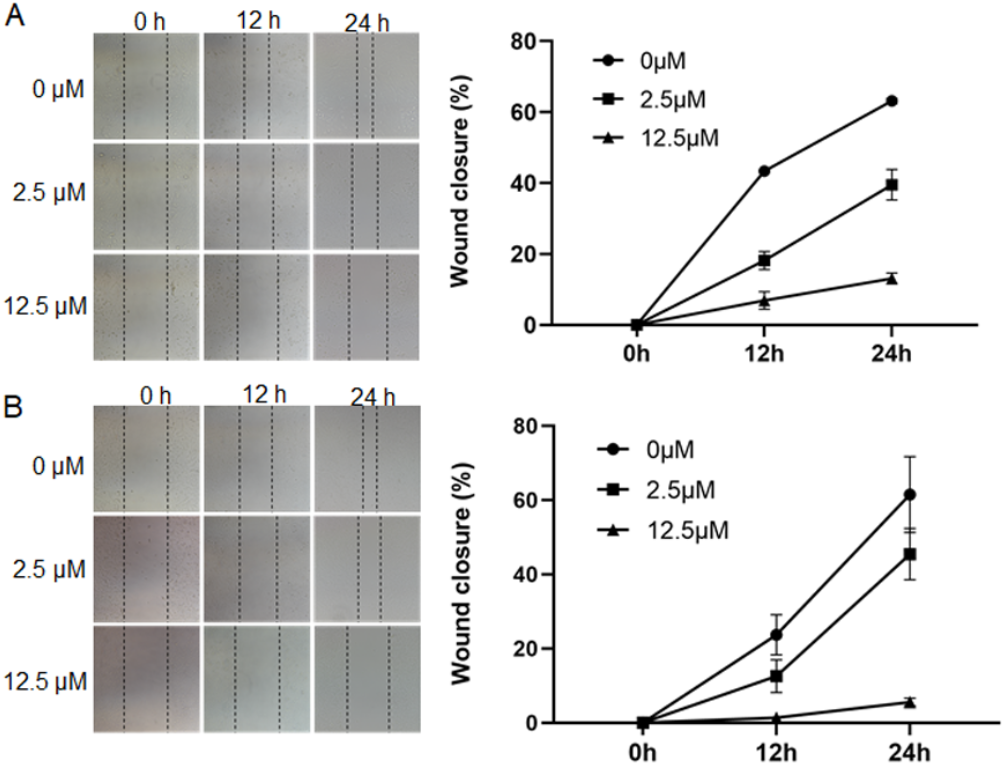
Wound healing assay. Compound AK-778-XXMU suppresses glioma cell migrate. After creating a straight physical “wound”, GL261 and U87 were treated with different concentrations (0, 2.5, 12.5µM) of compound AK-778-XXMU for 24 h. The process of cells migration into the gap were monitored by taking typical photos at different time points (0, 12 and 24 h). Data points represent the mean ± SD of three independent experiments.

**Fig. 6.**
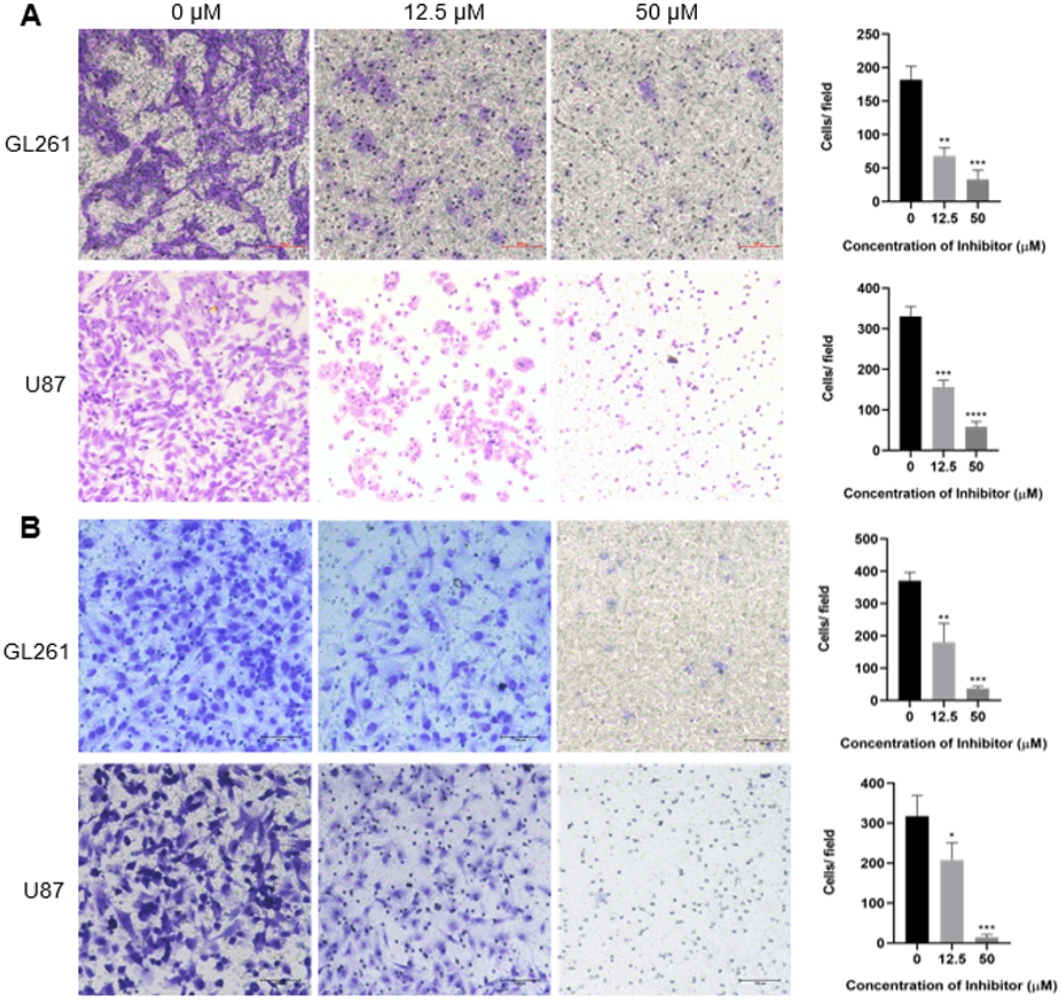
Transwell assay. A: representative images of migrated cells treated with different concentration of compound AK-778-XXMU and the quantitative analysis of migrated cells. B: representative images of invaded cells treated at different concentration of compound AK-778-XXMU and the quantitative analysis of invaded cells. X axis represents different concentrations of compound AK-778-XXMU treatment, and Y axis is the average number of migrated cells or invaded cells in three independent experiments at least. Each treatment group was compared with the control group. (\**p*<0.05; \*\**p*<0.01; \*\*\**p* <0.001; \*\*\*\**p* <0.0001 vs. Control).

### 6. Compound AK-778-XXMU induces apoptosis in glioma cells

To explore the effect of AK-778-XXMU on induction of apoptosis in glioma cells, Annexin V/PI double stained with flow cytometry was performed *in vitro*. Cells were treated with 25 µM and 50 µM AK-778-XXMU for 24 h. It was shown in **Fig. 7** that exposures of U87 and GL261 tumor cells to different concentrations of AK-778-XXMU resulted in increased apoptosis.

**Fig. 7.**
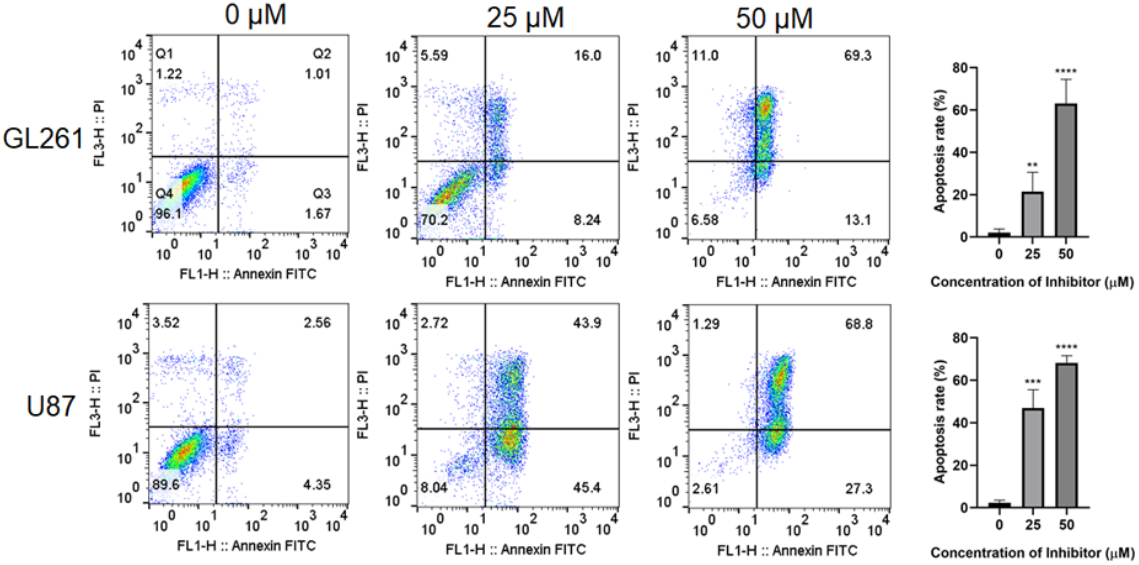
Compound AK-778-XXMU induces apoptosis in glioma cells. Human astroblastoma (U87) and mouse glioma (GL261) were treated at different concentrations (0, 25, 50 µM) of compound AK-778-XXMU for 24 h. Significantly increased apoptosis percentage of GL261 (A) and U87 (B) were detected by Annexin V/PI double stained by flow cytometry. Data represents the mean ± SD of at least three independent experiments. (\*\*\**p*<0.001 vs. Control; \*\*\*\**p*<0.0001 vs. Control).

### 7. Effect of compound AK-778-XXMU on ID2-KDR signaling

Glioma has abundant neovascularization network to support its invasive growth. KDR, also named VEGFR2, as an endotheliocyte-specific tyrosine kinase receptor, is critical in VEGF signaling and proangiogenic response in glioma malignant progression [43,44]. It has been reported that ID2 plays a critical role in KDR activation [17]. Therefore, western blot was performed to determine whether ID2-KDR signaling could be impaired by compound AK-778-XXMU. As expected, compared with the control group (0 µM), three proteins involved in ID2-KDR signaling (ID2, VEGFA, KDR) were greatly decreased in a dose-dependent manner (**Fig. 8**), and this expression pattern was validated by qRT-PCR (**Fig. 9**). Moreover, with increasing concentrations of compound AK-778-XXMU, an accumulation of tumor suppressor gene *PTEN* and an impaired expression of MMP2,MMP9 were found (**Fig. 8**).

**Fig. 8.**
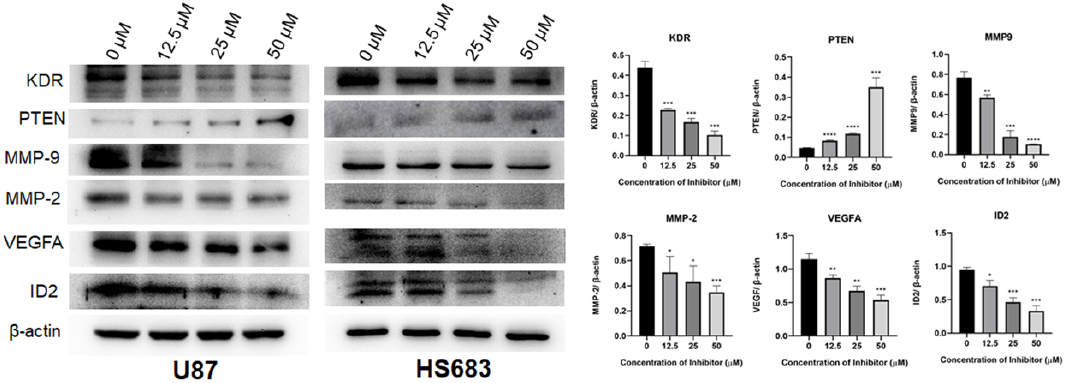
Compound AK-778-XXMU inhibits ID2-KDR signaling using Western blotting. Human glioma cells U87 and HS683 were treated at different concentrations of compound AK-778-XXMU for 24 h. The expression of ID2, VEGFA, PTEN, MMP2, MMP9 and KDR were detected in the whole cell lysates by Western blot. β-actin was used as the internal reference protein for each sample. The data were normalized to β-actin expression. (\**p*<0.05; \*\**p*<0.01; \*\*\**p*<0.001; \*\*\*\**p*<0.0001 vs. Control).

**Fig. 9.**
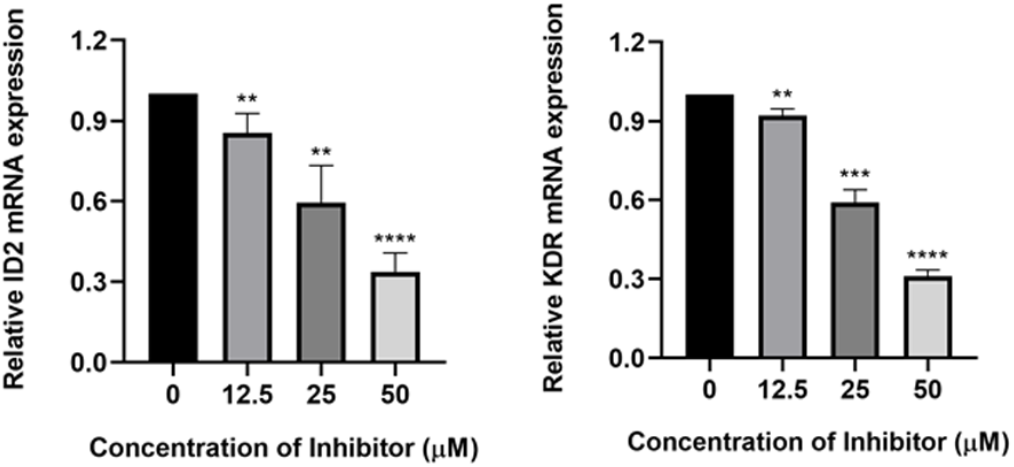
qRT-PCR analysis of ID2 and KDR mRNA expression level. The mRNA expression levels of ID2 and KDR (VEGFR2) in U87 under the same conditions were assessed by qRT-PCR. The data were normalized to GAPDH expression. (\*\**p*<0.01; \*\*\**p*<0.001; \*\*\*\**p* <0.0001 vs. Control)

### 8. Compound AK-778-XXMU slows down tumor growth *in vivo*

Inspired by the potent effect of AK-778-XXMU in inhibiting glioma cell growth *in vitro*, next we inoculated our NCG mouse with xenograft U87 cells into the right axilla to determine the antitumor effect of compound AK-778-XXMU *in vivo*. As shown in **Fig. 10**, in contrast to control group, tumors in treatment group (AK-778-XXMU: 2.5mg/kg) had significantly smaller volumes at the endpoint with 49.7% tumor inhibitory rate. Furthermore, the treatment group showed a reduction of tumor weight of 57.8% compared with control group at day 12. Notably, there was no significant reduction between the two groups in body weight. Overall, these data confirmed that compound AK-778-XXMU could inhibit tumor growth, possibly by impairing ID2 functions in glioma without obvious side-effects.

**Fig. 10.**
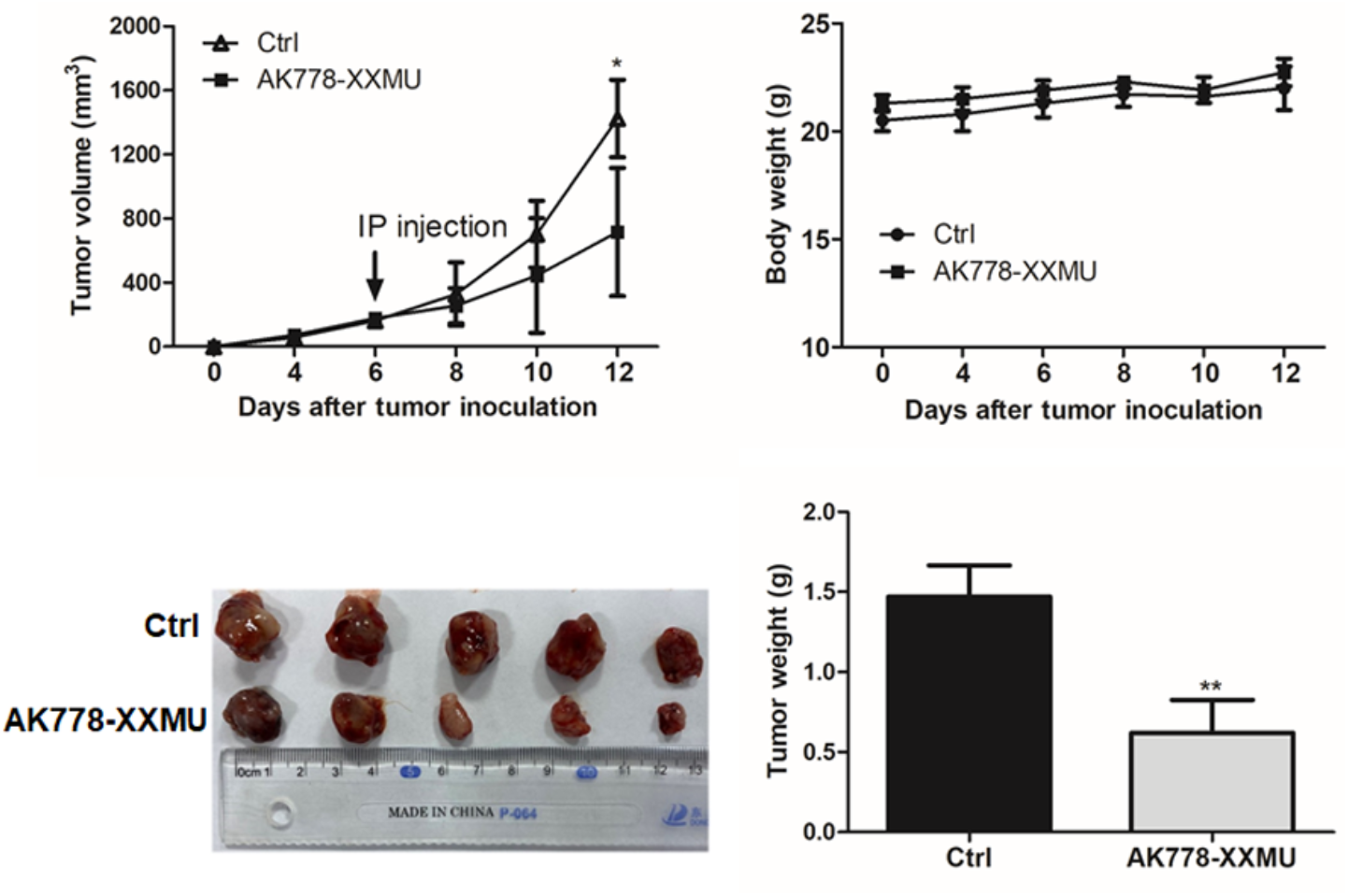
Inhibitory effects of compound AK-778-XXMU in U87 glioma in vivo. NCG mice bearing U87 tumors were injected with DMSO for control group or compound AK-778-XXMU for treatment group (2.5mg/kg every other day for 5 days, n=5 for each group). A: Tumor volume growth curve after compound AK-778-XXMU treatment. B: Body weight change of the mice. C and D: the excised tumor and the tumor weight graph at the end of experiment. (\*\**p*< 0.01 vs Control).

To provide more insight into the experimental results above that compound AK-778-XXMU could inhibit the growth of the tumor in NCG mice and further understand the differences of tumor growth between the two groups, we removed tumor blocks from the armpit of ten mice after euthanasia of them. As a proliferating cell associated nuclear antigen, Ki67 antigen was expressed in G1, S, G2 and M phases of cell cycle, but not in quiescent phase G0 [45]. Its function is closely related with cell proliferation, which means the higher positive ratio of Ki67, the faster tumor growth. A large number of neovascularization is a typical feature of malignant tumors and often an important cause of distant metastasis of tumors. A recent study demonstrated that glioblastoma stem cells could differentiate into endothelial cells which advance angiogenesis [46,47]. CD31 antigen, also known as platelet endothelial cell adhesion molecule-1 (PECAM-1), is a classical marker of endothelial cell that is also strongly related to the malignant progression of glioma [46,48]. Then immunohistochemical staining was performed on the interior of the tumor. As shown in **Fig. 11A** and **11B**, after treated with compound AK-778-XXMU, notable reduced positive rates of Ki67 and CD31 were detected inside the tumors with no destructive side effects on normal tissues (**Fig. 11C**) by staining the heart, liver, lungs, and kidneys with hematoxylin & eosin (HE) in treatment group, further demonstrating that compound AK-778-XXMU had a potent inhibitory effect on glioma.

**Fig. 11.**
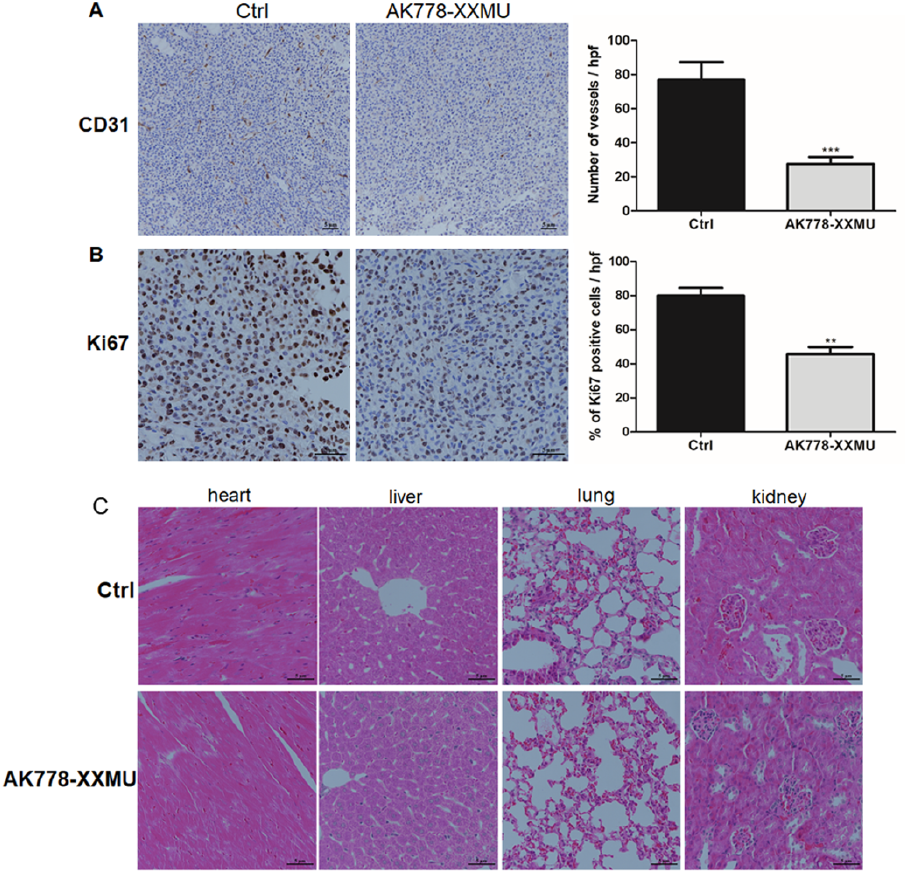
Immunohistological anlaysis. Representative immunohistochemical staining photographs of endothelial marker CD31 (A) for tumors vessels and proliferation marker Ki67 (left B) for cell proliferation in U87 tumors from NCG tumor-bearing mice at endpoint. (C) Histological analysis of different tissues from tumor-bearing NCG mice at endpoint. Different tissues (heart, liver, lung, kidney) from U87 tumor-bearing mice were fixed with 10 % (v/v) formalin and stained using Hematoxylin and Eosin (H&E). No histological and morphological heterogeneity of these viscera from tumor-bearing NCG mice in two groups were detected.

## Conclusions

Given the poor prognosis of glioma patients, it has become imperative to search for new targeted therapeutic regimen to control the malignant progression of glioma and improve the life quality of patients. In view of the strong correlation between ID2 protein and glioma, a pharmacophore-based virtual screening was conducted to obtain 16 small-molecule compounds, of which 13 had IC_50_ values lower than 100 μM from CCK8 assays. Further studies were focused on compound AK-778-XXMU, which is an indole derivative. The affinity between AK-778-XXMU and ID2 was determined to be 129 nM by SPR experiment, which is in the low submicromolar range. Its binding mode was studied by comparison with AGX51 in the binding site of ID2 protein, providing the explanation of its potency at the atomic level. It was also shown that AK-778-XXMU could inhibit cell migration and invasion of glioma cell lines, induce apoptosis, and more importantly, slow down the tumor growth *in vivo*. Therefore, compound AK-778-XXMU could be a promising agent for the treatment of glioma with ID2 overexpression in the future.

## Material and methods

### 1. Cell lines and cell culture

Cell lines (Human astroblastoma, U87; human glioma, HS683; mouse glioma, GL261) used in this study were obtained from ATCC. All these cell lines were cultured in Dulbecco’s Modified Eagle’s medium (DMEM High Glucose; Hyclone, USA) containing 10% fetal bovine serum (FBS) (Gibco, New York, USA), 1% (v/v) antibiotic solution containing penicillin and streptomycin (Sigma-Aldrich, USA) in a 5% CO_2_ atmosphere at 37 °C.

### 2. Molecular modeling and virtual screening

Compound AGX51 was used to build the pharmacophores. The 3D model of AGX51 was built and fully optimized with Discovery Studio 3.5 (BIOVIA, Dassault Systèmes). Since AGX51 has one chiral center, both enantiomers were used and two pharmacophores were generated by LigandScout 4.0 software by Inte:Ligand GmbH (http://www.inteligand.com/ligandscout/) [40]. They were subjected to commercial the following databases: CHEMBRIDGE, SPECS, CHEMDIV, ENAMINE, IBS, LIFECHEMICALS, VITAS-M (2,281 compounds in total) provided in their electronic form by Shanghai-based Topscience Co., Ldt (https://www.tsbiochem.com/) for virtual screening also in LigandScout. The pharmacophore models built from the *R* and *S* configurations yielded 24 and 33 hits, respectively. In the end, 16 compounds were purchased after consideration of possible future chemical modifications. Molecular docking was performed using Glide from Schrödinger, Inc using the default XP protocol [49,50].

### 3. CCK8 assay

CCK8 assay was used to elucidate the cytotoxicity of inhibitor on glioma cells. The cells were seeded into 96-well plate at 4 × 10^3^ cells per well with 100 µL DMEM supplemented with 10% fetal bovine serum and incubated at 37 °C. When the cells were grown to 60-70% confluency, different concentrations (0, 1.56, 3.125, 6.25, 12.5, 25, 50 µM) of inhibitor were added. After incubation for 24 h, 10 µL CCK-8 was added to each well. The absorbance of each well was detected at 450 nm after incubating at 37 °C for 4 hours. Every experiment was performed in triplicates.

### 4. SPR

In SPR assay, the surface of the Series S Sensor CM5 chip (GE, BR-1005-30) was activated with 400 mM EDC and 100 mM NHS at a flow rate of 10 μL/min for 420 s. Then human ID2 protein (Cat#PROTQ02363, purchased from Boster Biological Technology Co. Ltd, China.) was diluted to 50 μg/mL with fixative reagent (10 mM sodium acetate, pH 4.5) and immobilized on the surface of Chip CM5 with a fixed volume of approximately 6000 RU prior to the measurement of the compound AK-778-XXMU interaction. For compounds, they were first diluted 20-fold with dilution buffer (1 × PBS, 0.05% Tween20) to change the DMSO content to 5%, and then diluted with running reagent (1 × PBS, 0.05% Tween20, 5% DMSO) at concentrations of 100, 50, 25, 12.5, 6.25, 3.125, and 0 μM, respectively. The diluted compounds were injected into the experimental channel and the reference channel at a flow rate of 30 μL/min with a binding time of 60 s and a dissociation time of 120 s. The KD value of each antibody was calculated using Biacore 8K (GE Healthcare) analysis software. All SPR measurements were performed in triplicates.

### 5. Apoptosis analysis

Apoptosis of U87 and GL261 cells were detected by Annexin V-FITC/PI apoptosis detection kit (Key Gen Bio TECH). Glioma cells were treated with different concentrations (0, 25, 50 µM) of inhibitors for 24 h, harvested and washed twice with PBS and centrifuged. All cells were then suspended in 500 μL binding buffer, and stained with 5 μL Annexin V–FITC and 5 μL PI for 20 min at room temperature in the dark. Samples were run on a flow cytometer within 1 h.

### 6. Wound healing assay

In the migration assay, glioma cells were planted into 6-well plate at the density of 5 × 105 cells per well with 2 mL high-glucose DMEM containing 10% FBS, incubated at 37 °C in a 5% CO_2_ incubator for 24 h. Then a straight physical “wound” within cell monolayer was created. The cells were washed twice with PBS buffer gently and the cells that have been scratched out were discarded. 2 mL high glucose medium (DMEM) containing different concentrations of compound (0 μM, 2.5 μM, 12.5 μM) was added to each well. Then continue to culture in 37 °C incubator. Monitor the process of cell migration into the gap by taking photos at different time points (0 h, 12 h, 24 h).

### 7. Migration assay

In the migration assay, Transwell inserts with an 8 µM pore size were placed in 24-well plates with 500 µL DMEM culture medium added 10% fetal bovine serum. The U87 and HS683 cells were diluted to 2×10^5^ cells/mL with DMEM high glucose medium with 2% fetal bovine serum. A total of 200 µL of cell suspension was added to the inserts. Meanwhile different concentrations (0, 12.5, 50 µM) of inhibitors were added. Then the chambers were cultured in a 5% CO_2_ incubator at 37°C for 24 h. The cells were fixed with 4% paraformaldehyde, stained with crystal violet then take pictures and count in five different fields with an inverted microscope (200×).

### 8. Invasion assay

In the invasion assay, 30 μL Matrigel (BD Bioscience) was added to the up chamber then allowed to polymerize for at least 40 min at 37°C. The U87 and HS683 cells were resuspended to 2×10^5^ cells/mL with DMEM high glucose medium with 2% fetal bovine serum. A total of 200 µL of cell suspension was added to the inserts. Meanwhile different concentrations (0, 12.5, 50µM) of inhibitors were added. Then the plates were cultured in a 5% CO_2_ incubator at 37 °C for 24 h. Gently wipe off the Matrigel (Corning, Cat#356234) in the upper chamber with a cotton swab. Then the cells were fixed with 4% paraformaldehyde, stained with crystal violet (Beyotime) then take pictures and count in five different fields with an inverted microscope Pierce (200×).

### 9. Western blotting

Glioma cells were treated with different concentrations (0, 12.5, 25, 50 µM) of compound for 24 hours before being lysed by RIPA containing protease inhibitor. Lysates were incubated on ice for 20 min prior to centrifugation at 12000 rpm for 20 min at 4°C. After being determined concentrations using a BCA kit (Pierce), 30 µg total protein was separated by SDS-PAGE then transferred to the polyvinylidene fluoride membrane. The primary antibodies used were as follows: anti-KDR (Cell Signaling Technology, #2479), anti-PTEN (Cell Signaling Technology, #9188), anti-MMP-9 (Cell Signaling Technology, #13667), anti-MMP-2 (Cell Signaling Technology, #87809), anti-VEGFA (Cell Signaling Technology, #65373), anti-ID2 (Cell Signaling Technology, #3431), anti-β-actin (Cell Signaling Technology, #4970). The dilutions for the antibodies were as follows: 1:1000 for KDR, 1:1000 for PTEN, 1:1000 for MMP-9, 1:1000 for MMP-2, 1:1000 for VEGFA, 1:1000 for ID2, 1:1000 for β-actin, 1:10000 for Goat anti-Rabbit IgG.

### 10. Real-time PCR

To quantify the genes involved ID2-KDR signaling expression, human glioma cells HS683 were exposed to different concentrations (0, 12.5, 25, 50µM) of the compound, after 24 h, cells were lysed by RNAiso Plus (Takara, 9108, Japan). Before performing RT-PCR, Prime Script RT reagent Kit (Takara, RR047A, Japan) were used to reverse transcribe the extracted RNA into cDNA according to the given instructions. The results were assessed by calculating 2^-ΔΔCt^. The sequences of primers are as follows: GAPDH forward primer: 5′-GGAGCGAGATCCCTCCAAAAT-3′, reverse primer: 5′-GGCTGTTGTCATACTTCTCATGG-3′; ID2 forward primer: 5′-AGTCCCGTGAGGTCCGTTAG-3′, reverse primer: 5′-AGTCGTTCATGTTGTATAGCAGG-3′; VEGFR2 forward primer: 5′-GTGATCGGAAATGACACTGGAG-3′, reverse primer: 5′-CATGTTGGTCACTAACAGAAGCA-3′.

### 11. Subcutaneous tumorigenesis in NCG mice

To generate mouse models, ten 8-week-old NCG mice were injected i.h. with human glioma cell U87 into the right axilla according to the number of 1 × 10^6^ cells/mouse. When the tumor volume was visible and reached 100 mm^3^, ten mice were randomly divided into two cohorts: control group (n=5) and treatment group (n=5). The control group (DMSO, 2.5 ml/kg), AK-778-XXMU (2.5mg/kg) compound was injected i.p. into the two groups tumor-bearing mice respectively. Control group received DMSO (1% v/v) on the 1st, 3rd and 5th day. Treatment group were injected with AK-778-XXMU compound (1% v/v, dissolved in DMSO) on the 1st, 3rd and 5th day. The body weight and tumor size of mice were measured at the same time of each administration, and the changes of body weight and activity ability of mice were dynamically monitored. On the endpoint, the 7th day of the first administration, the mice were euthanized, the tumor were excised and weighed.

### 12. IHC (Immunohistochemistry) assay

The expression of micro vessel density marker CD31 and cell proliferation index Ki67 in tumor samples from tumor-bearing NCG mice were tested via IHC. IHC staining was performed as previously described [51]. Each slice was taken five views for photographing at 400 X magnification for Ki67 and at 200 X magnification for CD31. Each obvious area of positive staining for CD31 was counted as a single vessel. Results were expressed as mean number of Ki67 positive cells ± SEM per high-power field (X400) or the mean number of micro vessels for CD31 ± SEM per high-power field (X200). A total of 15 high-power fields was examined and the number of micro vessels for CD31 or Ki67 positive-staining cells in each group was averaged and expressed as the number per high power field.

### 13. Hematoxylin & eosin (HE) staining

After being euthanized, different tissues (heart, liver, lung, kidney) from ten tumor-bearing NCG mice were fixed with 10% (v/v) formalin over 24 hours. Then all the specimens were washed, dehydrated and embedded with paraffin commonly. 5µm-thick viscera sections were cut randomly, fixed onto the glass slides then, stained using Hematoxylin and Eosin (H&E) (Solarbio, G1121) after being dewaxed to water, the images were taken by light microscopy with Nikon software (Melville, NY, USA) at 400 × magnification at last.

### 14. Statistical analysis

All measurements were performed in triplicates. The quantitative data were performed with the SPSS 20.0 software package expressed as the means ± standard deviation (SD). The significance of the difference from the controls for each experimental test condition was assayed using unpaired t-test. ANOVAs with the post hoc Dunnett’s test were used for tumor growth in comparison to the control. *P* values < 0.05 were considered statistically significant.

## Supporting information

Biacore assay

purchased compounds list

## Declaration of Competing Interest

The authors declare that they have no known competing financial interests or personal relationships that could have appeared to influence the work reported in this paper.

## Acknowledgments

This work was supported by National Natural Science Foundation of China (No. 21406143) to Bin Lin, NSFC-Henan Union grant (No. U1904131) to Genshen Zhong. Bin Lin also would like to acknowledge the faculty startup funding from Shenyang Pharmaceutical University.

---

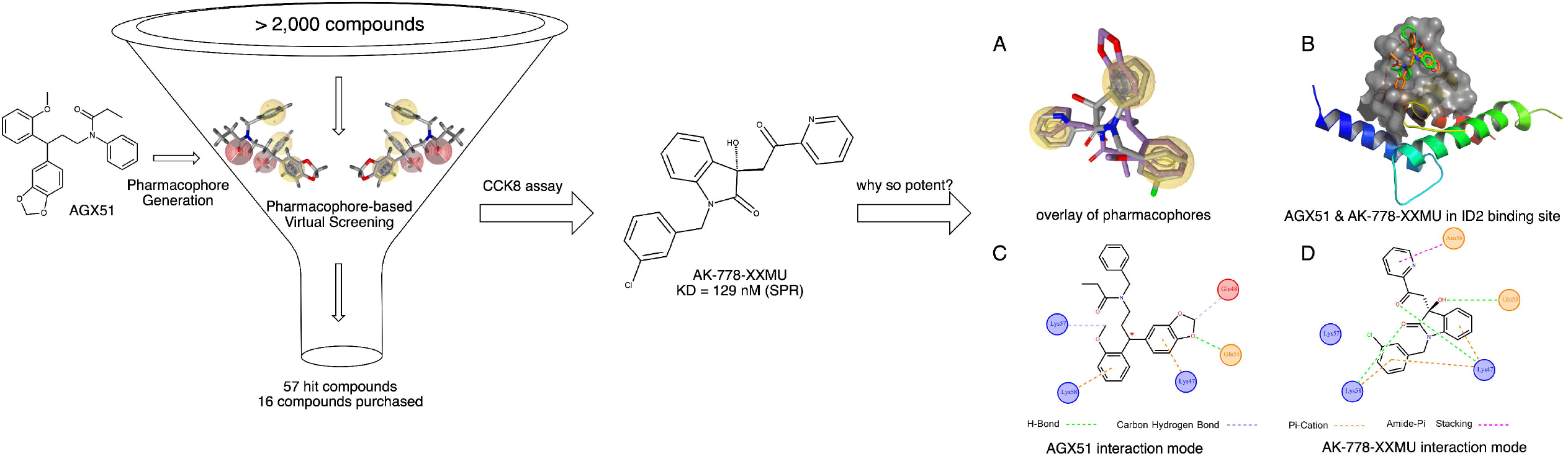

## Notes

### Competing Interest Statement

The authors have declared no competing interest.

### Summary of Updates

One of the authors, Yujie Huang, wished to withdraw from the publication. His name has been removed from this version.

## References

[1] E.G. Van Meir, C.G. Hadjipanayis, A.D. Norden, H.K. Shu, P.Y. Wen, J.J. Olson, Exciting new advances in neuro-oncology: the avenue to a cure for malignant glioma, CA. Cancer J. Clin. 60 (2010) 166–193. https://doi.org/10.3322/caac.20069.

[2] A. Omuro, L.M. DeAngelis, Glioblastoma and other malignant gliomas: a clinical review, JAMA. 310 (2013) 1842–1850. https://doi.org/10.1001/jama.2013.280319.

[3] T.A. Dolecek, J.M. Propp, N.E. Stroup, C. Kruchko, CBTRUS statistical report: primary brain and central nervous system tumors diagnosed in the United States in 2005-2009, Neuro-Oncol. 14 (2012) v1–49. https://doi.org/10.1093/neuonc/nos218.

[4] P.Y. Wen, E.Q. Lee, D.A. Reardon, K.L. Ligon, W.K.A. Yung, Current clinical development of PI3K pathway inhibitors in glioblastoma, Neuro-Oncol. 14 (2012) 819–829. https://doi.org/10.1093/neuonc/nos117.

[5] H. Malkki, Neuro-oncology: bevacizumab prolongs progression-free survival but not overall survival in newly diagnosed glioblastoma, Nat. Rev. Neurol. 10 (2014) 179. https://doi.org/10.1038/nrneurol.2014.47.

[6] S. Bao, Q. Wu, R.E. McLendon, Y. Hao, Q. Shi, A.B. Hjelmeland, M.W. Dewhirst, D.D. Bigner, J.N. Rich, Glioma stem cells promote radioresistance by preferential activation of the DNA damage response, Nature 444 (2006) 756–760. https://doi.org/10.1038/nature05236.

[7] I. Paw, R.C. Carpenter, K. Watabe, W. Debinski, H.W. Lo, Mechanisms regulating glioma invasion, Cancer Lett. 362 (2015) 1–7. https://doi.org/10.1016/j.canlet.2015.03.015.

[8] L.C. Hou, A. Veeravagu, A.R. Hsu, V.C. Tse, Recurrent glioblastoma multiforme: a review of natural history and management options, Neurosurg. Focus. 20 (2006) E5. https://doi.org/10.3171/foc.2006.20.4.2.

[9] R. Benezra, R.L. Davis, D. Lockshon, D.L. Turner, H. Weintraub, The protein Id: a negative regulator of helix-loop-helix DNA binding proteins, Cell. 61 (1990) 49–59. https://doi.org/10.1016/0092-8674(90)90214-y.

[10] H.A. Sikder, M.K. Devlin, S. Dunlap, B. Ryu, R.M. Alani, Id proteins in cell growth and tumorigenesis, Cancer Cell. 3 (2003) 525–530. https://doi.org/10.1016/s1535-6108(03)00141-7.

[11] X.H. Sun, N.G. Copeland, N.A. Jenkins, D. Baltimore, Id proteins Id1 and Id2 selectively inhibit DNA binding by one class of helix-loop-helix proteins, Mol. Cell. Biol. 11 (1991) 5603–5611. https://doi.org/10.1128/mcb.11.11.5603.

[12] J. Biggs, E. V Murphy, M.A. Israel, A human Id-like helix-loop-helix protein expressed during early development, Proc. Natl. Acad. Sci. 89 (1992) 1512–1516. https://doi.org/10.1073/pnas.89.4.1512.

[13] V. Riechmann, I. van Crüchten, F. Sablitzky, The expression pattern of Id4, a novel dominant negative helix-loop-helix protein, is distinct from Id1, Id2 and Id3, Nucleic Acids Res. 22 (1994) 749–755. https://doi.org/10.1093/nar/22.5.749.

[14] B.A. Christy, L.K. Sanders, L.F. Lau, N.G. Copeland, N.A. Jenkins, D. Nathans, An Id-related helix-loop-helix protein encoded by a growth factor-inducible gene, Proc. Natl. Acad. Sci. 88 (1991) 1815–1819. https://doi.org/10.1073/pnas.88.5.1815.

[15] J. Perk, A. Iavarone, R. Benezra, Id family of helix-loop-helix proteins in cancer, Nat. Rev. Cancer. 5 (2005) 603–614. https://doi.org/10.1038/nrc1673.

[16] Z. Dong, S. Liu, C. Zhou, T. sumida, H. Hamakawa, Z. Chen, P. Liu, F. Wei, Overexpression of Id-1 is associated with tumor angiogenesis and poor clinical outcome in oral squamous cell carcinoma, Oral Oncol. 46 (2010) 154–157. https://doi.org/10.1016/j.oraloncology.2009.11.005.

[17] Y. Huang, P. Rajappa, W. Hu, C. Hoffman, B. Cisse, J.H. Kim, E. Gorge, R. Yanowitch, W. Cope, E. Vartanian, R. Xu, T. Zhang, D. Pisapia, J. Xiang, J. Huse, I. Matei, H. Peinado, J. Bromberg, E. Holland, B.S. Ding, S. Rafii, D. Lyden, J. Greenfield, A proangiogenic signaling axis in myeloid cells promotes malignant progression of glioma, J. Clin. Invest. 127 (2017) 1826–1838. https://doi.org/10.1172/JCI86443.

[18] D.A. Vandeputte, D. Troost, S. Leenstra, H. Ijlst-Keizers, M. Ramkema, D.A. Bosch, F. Baas, N.K. Das, E. Aronica, Expression and distribution of id helix-loop-helix proteins in human astrocytic tumors, Glia. 38 (2002) 329–338. https://doi.org/10.1002/glia.10076.

[19] Z. Zhang, G.J. Rahme, P.D. Chatterjee, M.C. Havrda, M.A. Israel, ID2 promotes survival of glioblastoma cells during metabolic stress by regulating mitochondrial function, Cell Death Dis. 8 (2017) e2615. https://doi.org/10.1038/cddis.2017.14.

[20] A.J. Levine, A.M. Puzio-Kuter, The control of the metabolic switch in cancers by oncogenes and tumor suppressor genes, Science 330 (2010) 1340–1344. https://doi.org/10.1126/science.1193494.

[21] C. Stockmann, A. Doedens, A. Weidemann, N. Zhang, N. Takeda, J.I. Greenberg, D.A. Cheresh, R.S. Johnson, Deletion of vascular endothelial growth factor in myeloid cells accelerates tumorigenesis, Nature 456 (2008) 814–818. https://doi.org/10.1038/nature07445.

[22] R. Chen, M.C. Nishimura, S.M. Bumbaca, S. Kharbanda, W.F. Forrest, I.M. Kasman, J.M. Greve, R.H. Soriano, L.L. Gilmour, C.S. Rivers, Z. Modrusan, S. Nacu, S. Guerrero, K.A. Edgar, J.J. Wallin, K. Lamszus, M. Westphal, S. Heim, James, S.R. VandenBerg, J.F. Costello, S. Moorefield, C.J. Cowdrey, M. Prados, H.S. Phillips, A hierarchy of self-renewing tumor-initiating cell types in glioblastoma, Cancer Cell 17 (2010) 362–375. https://doi.org/10.1016/j.ccr.2009.12.049.

[23] S.M. Pollard, K. Yoshikawa, I.D. Clarke, D. Danovi, S. Stricker, R. Russell, J. Bayani, R. Head, M. Lee, M. Bernstein, J.A. Squire, A. Smith, P. Dirks, Glioma stem cell lines expanded in adherent culture have tumor-specific phenotypes and are suitable for chemical and genetic screens, Cell Stem Cell 4 (2009) 568–580. https://doi.org/10.1016/j.stem.2009.03.014.

[24] C. Roschger, C. Cabrele, The Id-protein family in developmental and cancer-associated pathways, Cell Commun Signal. 15 (2017) 7. https://doi.org/10.1186/s12964-016-0161-y.

[25] D. Lyden, A.Z. Young, D. Zagzag, W. Yan, W. Gerald, R. O’Reilly, B.L. Bader, R.O. Hynes, Y. Zhuang, K. Manova, R. Benezra, Id1 and Id3 are required for neurogenesis, angiogenesis and vascularization of tumour xenografts, Nature 401 (1999) 670–677. https://doi.org/10.1038/44334.

[26] M.B. Ruzinova, R.A. Schoer, W. Gerald, J.E. Egan, P.P. Pandolfi, S. Rafii, K. Manova, V. Mittal, R. Benezra, Effect of angiogenesis inhibition by Id loss and the contribution of bone-marrow-derived endothelial cells in spontaneous murine tumors, Cancer Cell 4 (2003) 277–289. https://doi.org/10.1016/s1535-6108(03)00240-x.

[27] A. Lasorella, R. Benezra, A. Iavarone, The ID proteins: master regulators of cancer stem cells and tumour aggressiveness, Nat. Rev. Cancer 14 (2014) 77–91. https://doi.org/10.1038/nrc3638.

[28] D. Gao, D.J. Nolan, A.S. Mellick, K. Bambino, K. McDonnell, V. Mittal, Endothelial progenitor cells control the angiogenic switch in mouse lung metastasis, Science 319 (2008) 195–198. https://doi.org/10.1126/science.1150224.

[29] J. Anido, A. Sáez-Borderías, A. Gonzàlez-Juncà, L. Rodón, G. Folch, M.A. Carmona, R.M. Prieto-Sánchez, I. Barba, E. Martínez-Sáez, L. Prudkin, I. Cuartas, C. Raventós, F. Martínez-Ricarte, M.A. Poca, D. García-Dorado, M.M. Lahn, J.M. Yingling, J. Rodón, J. Sahuquillo, J. Baselga, J. Seoane, TGF-β receptor inhibitors target the CD44high/Id1high glioma-initiating cell population in human glioblastoma, Cancer Cell 18 (2010) 655–668. https://doi.org/10.1016/j.ccr.2010.10.023.

[30] E. Henke, J. Perk, J. Vider, P. de Candia, Y. Chin, D.B. Solit, V. Ponomarev, L. Cartegni, K. Manova, N. Rosen, R. Benezra, Peptide-conjugated antisense oligonucleotides for targeted inhibition of a transcriptional regulator in vivo, Nat. Biotechnol. 26 (2008) 91–100. https://doi.org/10.1038/nbt1366.

[31] D.S. Mern, J. Hasskarl, B. Burwinkel, Inhibition of Id proteins by a peptide aptamer induces cell-cycle arrest and apoptosis in ovarian cancer cells, Br. J. Cancer. 103 (2010) 1237–1244. https://doi.org/10.1038/sj.bjc.6605897.

[32] H. Mistry, G. Hsieh, S.J. Buhrlage, M. Huang, E. Park, G.D. Cuny, I. Galinsky, R.M. Stone, N.S. Gray, A.D. D’Andrea, K. Parmar, Small-molecule inhibitors of USP1 target ID1 degradation in leukemic cells, Mol. Cancer Ther. 12 (2013) 2651–2662. https://doi.org/10.1158/1535-7163.Mct-13-0103-t.

[33] R. Murase, R. Kawamura, E. Singer, A. Pakdel, P. Sarma, J. Judkins, E. Elwakeel, S. Dayal, E. Martinez-Martinez, M. Amere, R. Gujjar, A. Mahadevan, P.-Y. Desprez, S.D. McAllister, Targeting multiple cannabinoid anti-tumour pathways with a resorcinol derivative leads to inhibition of advanced stages of breast cancer, Br. J. Pharmacol. 171 (2014) 4464–4477. https://doi.org/10.1111/bph.12803.

[34] L. Soroceanu, R. Murase, C. Limbad, E. Singer, J. Allison, I. Adrados, R. Kawamura, A. Pakdel, Y. Fukuyo, D. Nguyen, S. Khan, R. Arauz, G.L. Yount, D.H. Moore, P.-Y. Desprez, S.D. McAllister, Id-1 is a key transcriptional regulator of glioblastoma aggressiveness and a vovel therapeutic target, Cancer Res. 73 (2013) 1559–1569. https://doi.org/10.1158/0008-5472.Can-12-1943.

[35] P.M. Wojnarowicz, R. Lima e Silva, M. Ohnaka, S.B. Lee, Y. Chin, A. Kulukian, S.-H. Chang, B. Desai, M. Garcia Escolano, R. Shah, M. Garcia-Cao, S. Xu, R. Kadam, Y. Goldgur, M.A. Miller, O. Ouerfelli, G. Yang, T. Arakawa, S.K. Albanese, W.A. Garland, G. Stoller, J. Chaudhary, L. Norton, R.K. Soni, J. Philip, R.C. Hendrickson, A. Iavarone, A.J. Dannenberg, J.D. Chodera, N. Pavletich, A. Lasorella, P.A. Campochiaro, R. Benezra, A small-molecule pan-Id antagonist inhibits pathologic ocular neovascularization, Cell Rep. 29 (2019) 62–75. https://doi.org/10.1016/j.celrep.2019.08.073.

[36] M.V. Wong, S. Jiang, P. Palasingam, P.R. Kolatkar, A divalent ion is crucial in the structure and dominant-negative function of ID proteins, a class of helix-loop-helix transcription regulators, PLoS One. 7 (2012) e48591. https://doi.org/10.1371/journal.pone.0048591.

[37] S.-Y. Yang, Pharmacophore modeling and applications in drug discovery: challenges and recent advances, Drug Discov. Today. 15 (2010) 444–450. https://doi.org/10.1016/j.drudis.2010.03.013.

[38] G. Wolber, A.A. Dornhofer, T. Langer, Efficient overlay of small organic molecules using 3D pharmacophores, J. Comput. Aided. Mol. Des. 20 (2006) 773–788. https://doi.org/10.1007/s10822-006-9078-7.

[39] A.R. Leach, V.J. Gillet, R.A. Lewis, R. Taylor, Three-dimensional pharmacophore methods in drug discovery, J. Med. Chem. 53 (2010) 539–558. https://doi.org/10.1021/jm900817u.

[40] G. Wolber, T. Langer, LigandScout: 3-D pharmacophores derived from protein-bound ligands and their use as virtual screening filters, J. Chem. Inf. Model. 45 (2005) 160–169. https://doi.org/10.1021/ci049885e.

[41] C. Boozer, G. Kim, S. Cong, H. Guan, T. Londergan, Looking towards label-free biomolecular interaction analysis in a high-throughput format: a review of new surface plasmon resonance technologies, Curr. Opin. Biotechnol. 17 (2006) 400–405. https://doi.org/10.1016/j.copbio.2006.06.012.

[42] M.A. Cooper, Optical biosensors in drug discovery, Nat. Rev. Drug Discov. 1 (2002) 515–528. https://doi.org/10.1038/nrd838.

[43] A.K. Olsson, A. Dimberg, J. Kreuger, L. Claesson-Welsh, VEGF receptor signalling — in control of vascular function, Nat. Rev. Mol. Cell Biol. 7 (2006) 359–371. https://doi.org/10.1038/nrm1911.

[44] E. Krzywinska, C. Kantari-Mimoun, Y. Kerdiles, M. Sobecki, T. Isagawa, D. Gotthardt, M. Castells, J. Haubold, C. Millien, T. Viel, B. Tavitian, N. Takeda, J. Fandrey, E. Vivier, V. Sexl, C. Stockmann, Loss of HIF-1α in natural killer cells inhibits tumour growth by stimulating non-productive angiogenesis, Nat. Commun. 8 (2017) 1597. https://doi.org/10.1038/s41467-017-01599-w.

[45] M.J. Beresford, G.D. Wilson, A. Makris, Measuring proliferation in breast cancer: practicalities and applications, Breast Cancer Res. 8 (2006) 216. https://doi.org/10.1186/bcr1618.

[46] X. Mei, Y.S. Chen, F.R. Chen, S.Y. Xi, Z.P. Chen, Glioblastoma stem cell differentiation into endothelial cells evidenced through live-cell imaging, Neuro-Oncol. 19 (2017) 1109–1118. https://doi.org/10.1093/neuonc/nox016.

[47] D. Chroscinski, D. Sampey, N. Maherali, Reproducibility Project: Cancer Biology, Registered report: tumour vascularization via endothelial differentiation of glioblastoma stem-like cells, eLife. 4 (2015) e04363. https://doi.org/10.7554/eLife.04363.

[48] S. Das, P.A. Marsden, Angiogenesis in glioblastoma, N. Engl. J. Med. 369 (2013) 1561–1563. https://doi.org/10.1056/NEJMcibr1309402.

[49] R.A. Friesner, J.L. Banks, R.B. Murphy, T.A. Halgren, J.J. Klicic, D.T. Mainz, M.P. Repasky, E.H. Knoll, M. Shelley, J.K. Perry, D.E. Shaw, P. Francis, P.S. Shenkin, Glide: A new approach for rapid, accurate docking and scoring. 1. method and assessment of docking accuracy, J. Med. Chem. 47 (2004) 1739–1749. https://doi.org/10.1021/jm0306430.

[50] T.A. Halgren, R.B. Murphy, R.A. Friesner, H.S. Beard, L.L. Frye, W.T. Pollard, J.L. Banks, Glide: A new approach for rapid, accurate docking and scoring. 2. enrichment factors in database screening, J. Med. Chem. 47 (2004) 1750–1759. https://doi.org/10.1021/jm030644s.

[51] G. Zhong, S. Zhang, Y. Li, X. Liu, R. Gao, Q. Miao, Y. Zhen, A tandem scFv-based fusion protein and its enediyne-energized analogue show intensified therapeutic efficacy against lung carcinoma xenograft in athymic mice, xCancer Lett. 295 (2010) 124–133. https://doi.org/10.1016/j.canlet.2010.02.020.

