## supplementary material for "Discovery of novel ID2 antagonists from pharmacophore-based virtual screening as potential therapeutics for glioma"

The ^1^H-NMR and MS spectra of the purchase 16 compounds

^1^H-NMR of K784-9509

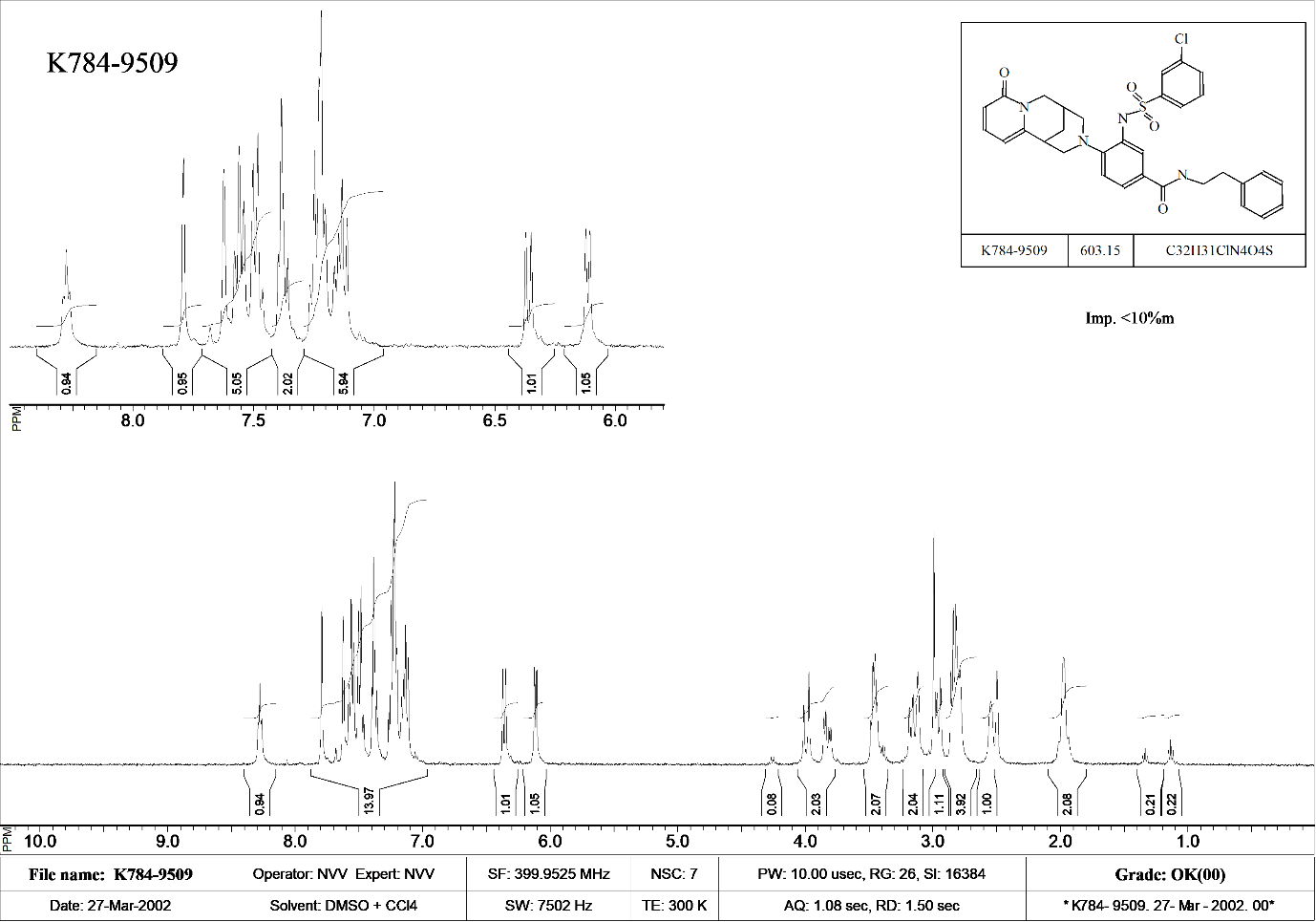

^1^H-NMR of C141-0292

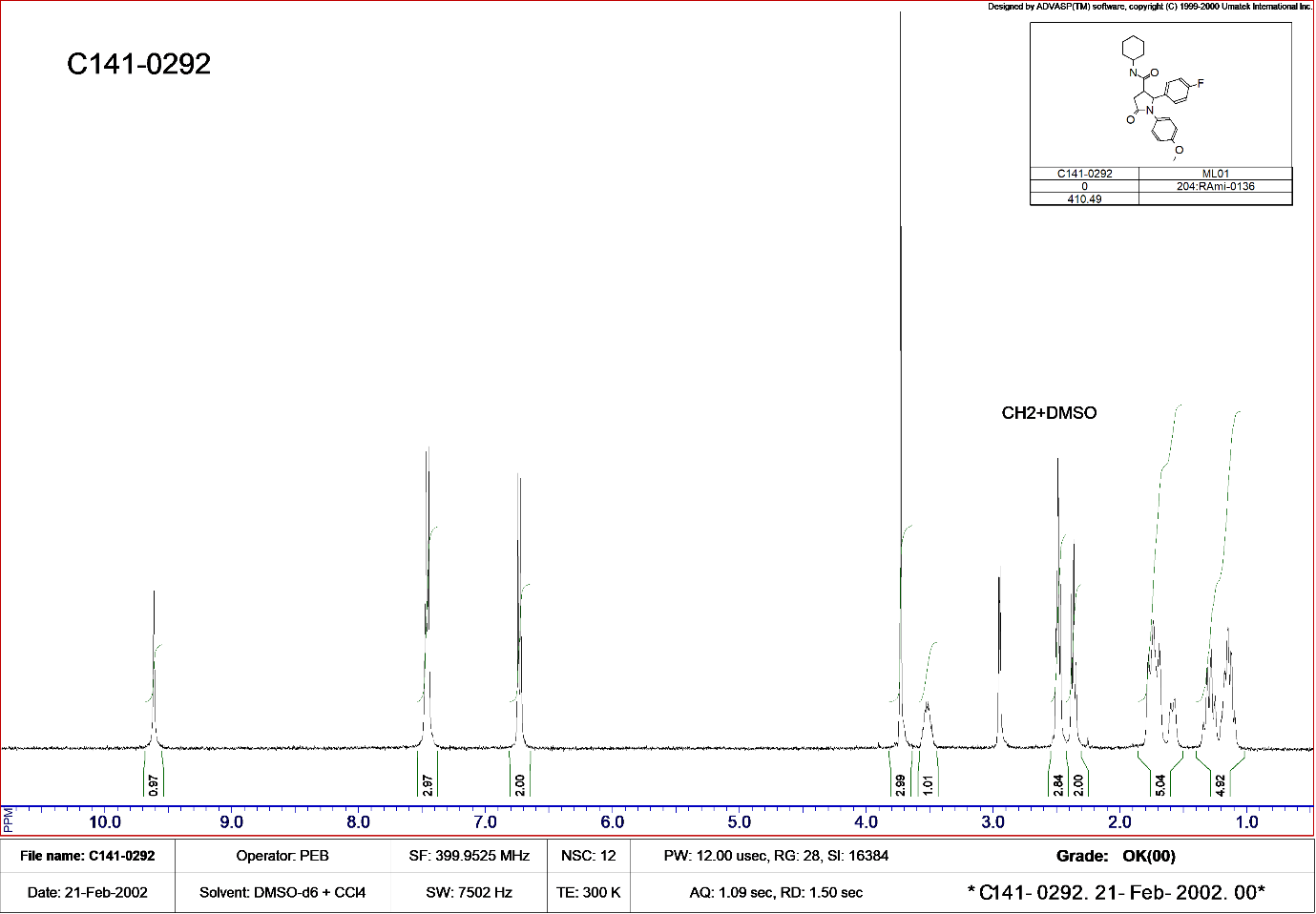

MS of AK-778/43465022

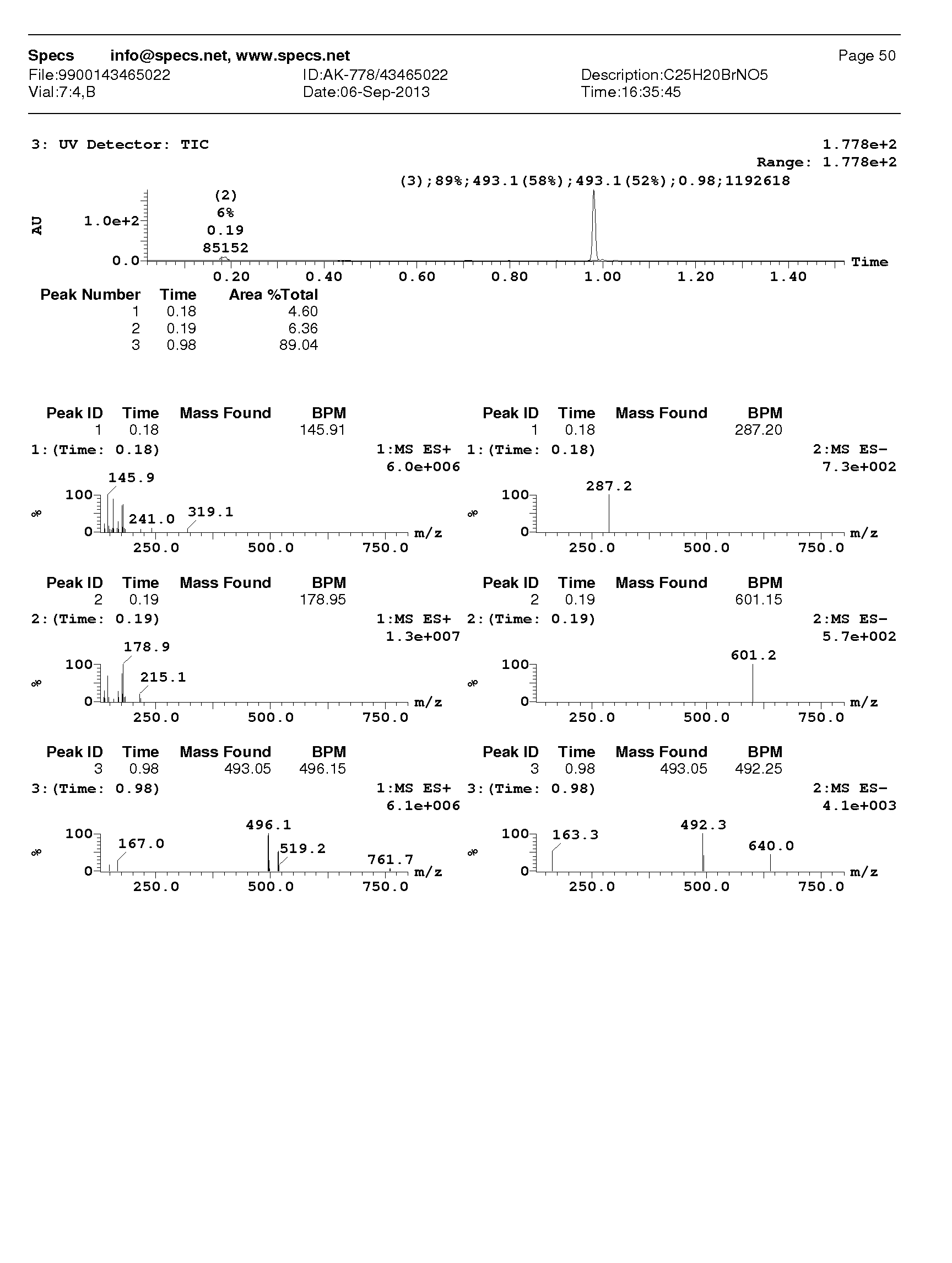

^1^H-NMR of C481-0437

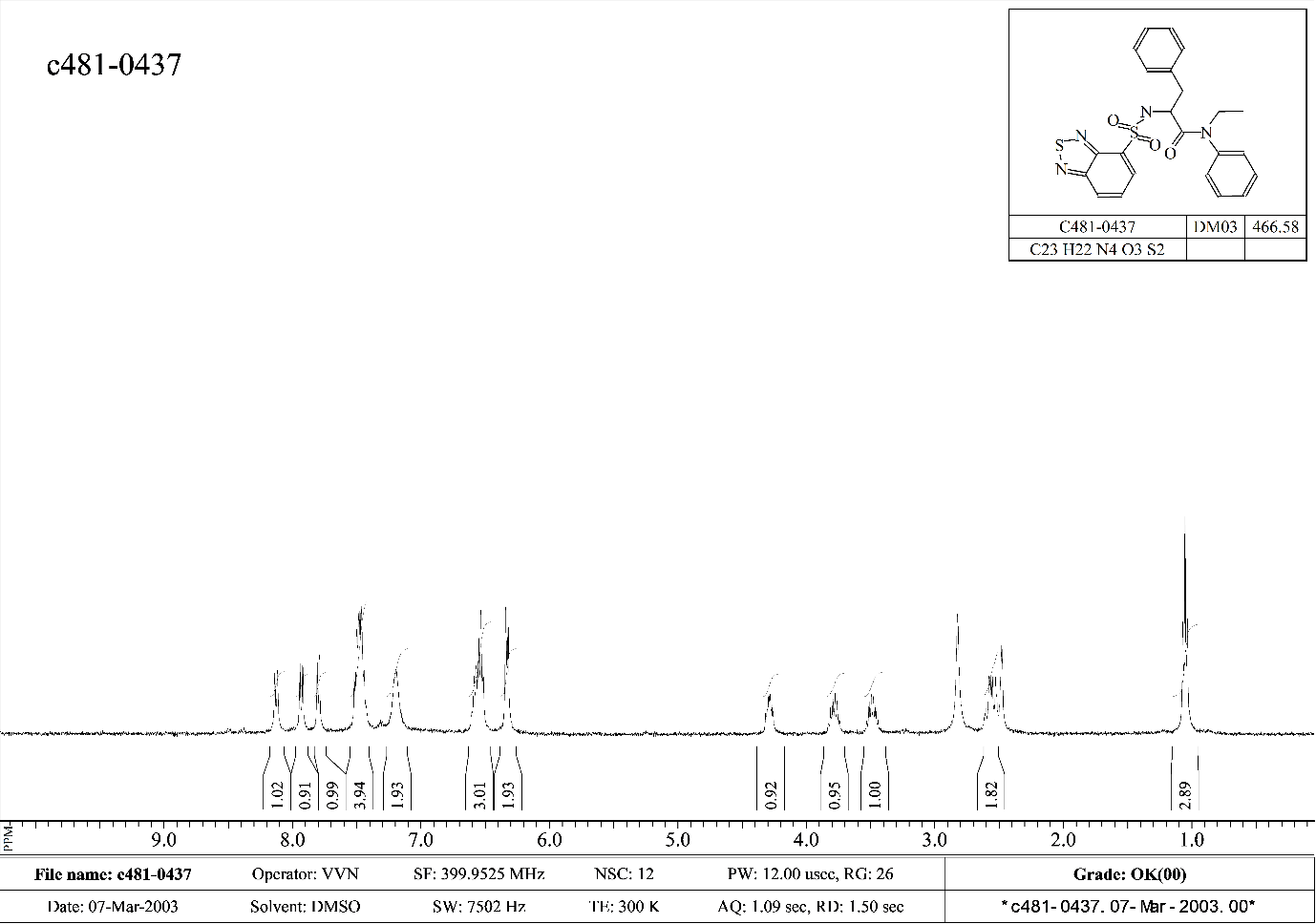

^1^H-NMR of 6835-3516

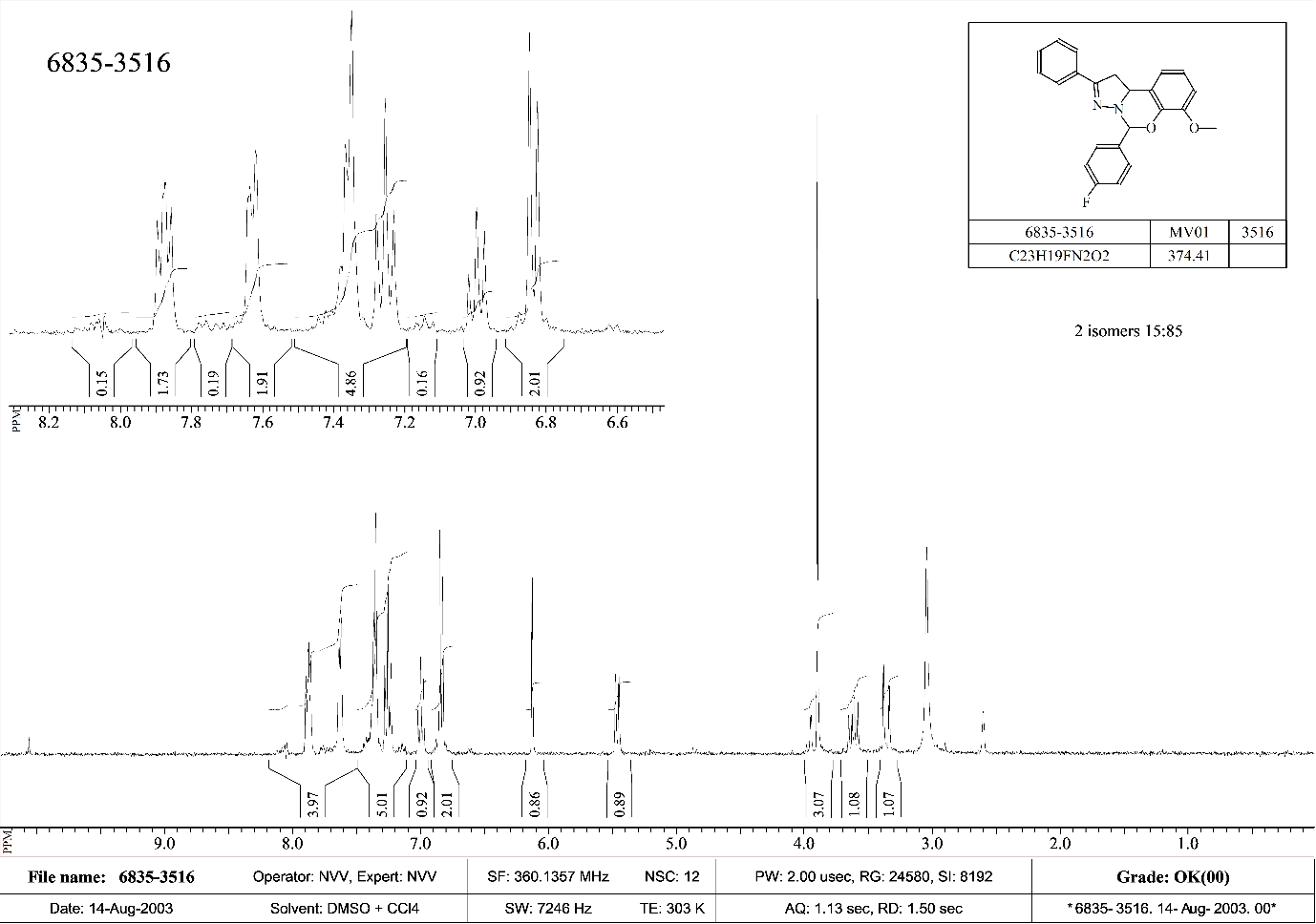

^1^H-NMR of E755-0111

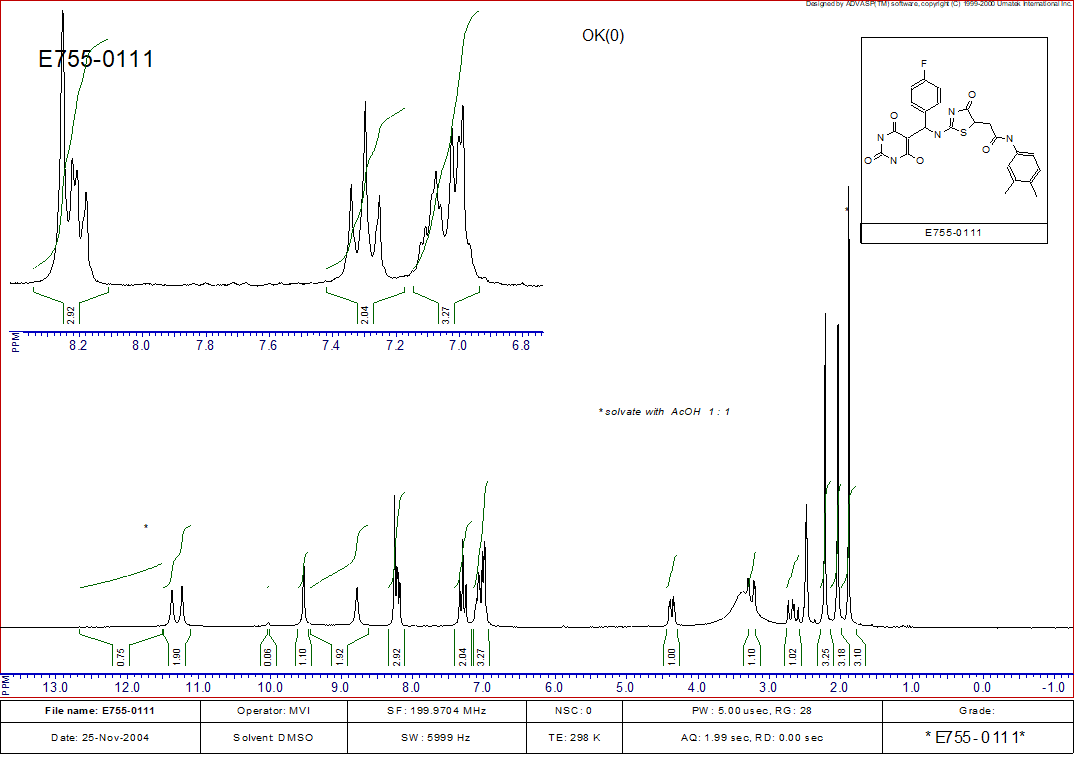

^1^H-NMR of Z32554393

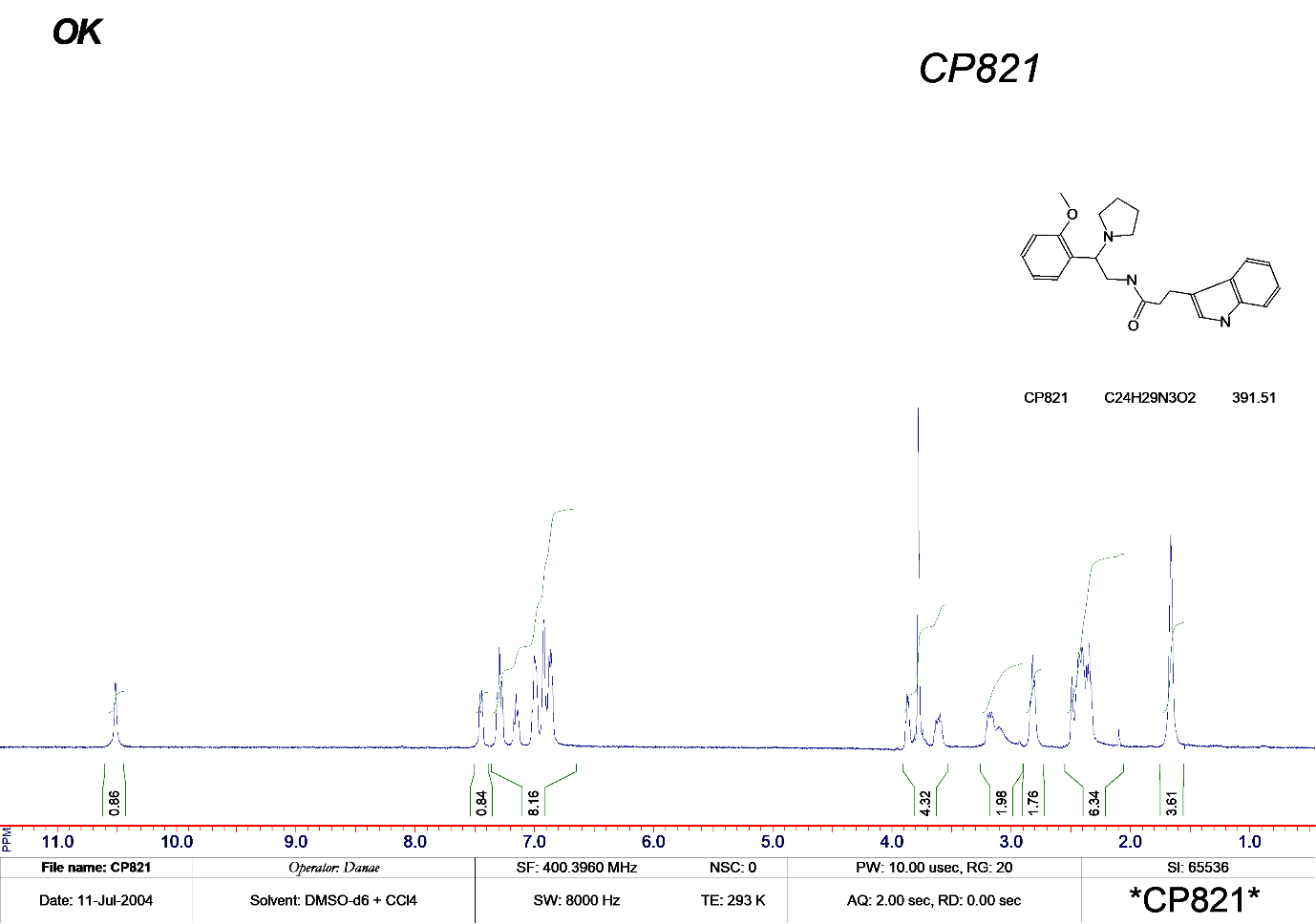

MS of Z32554393

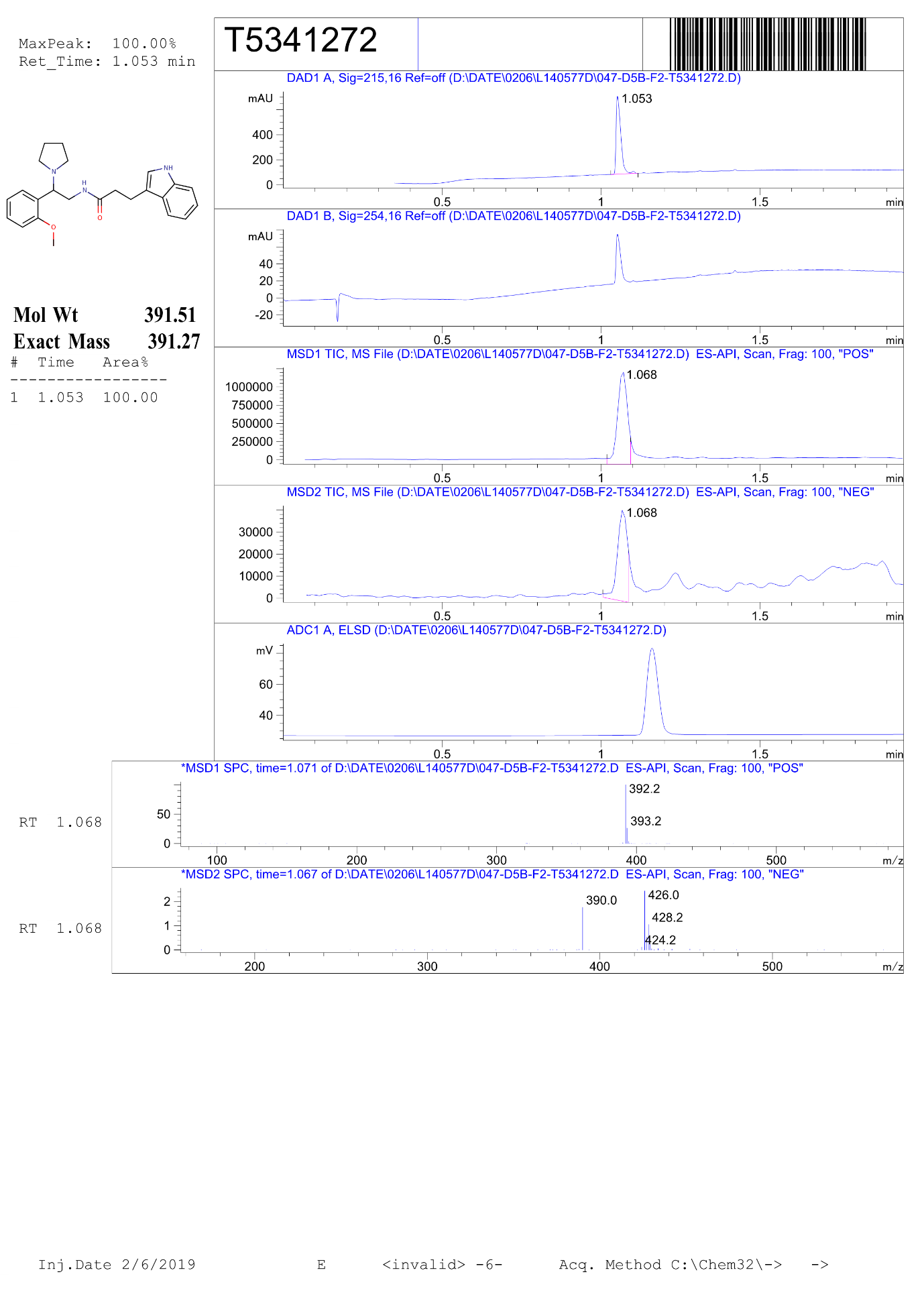

MS of AO-080/42479467

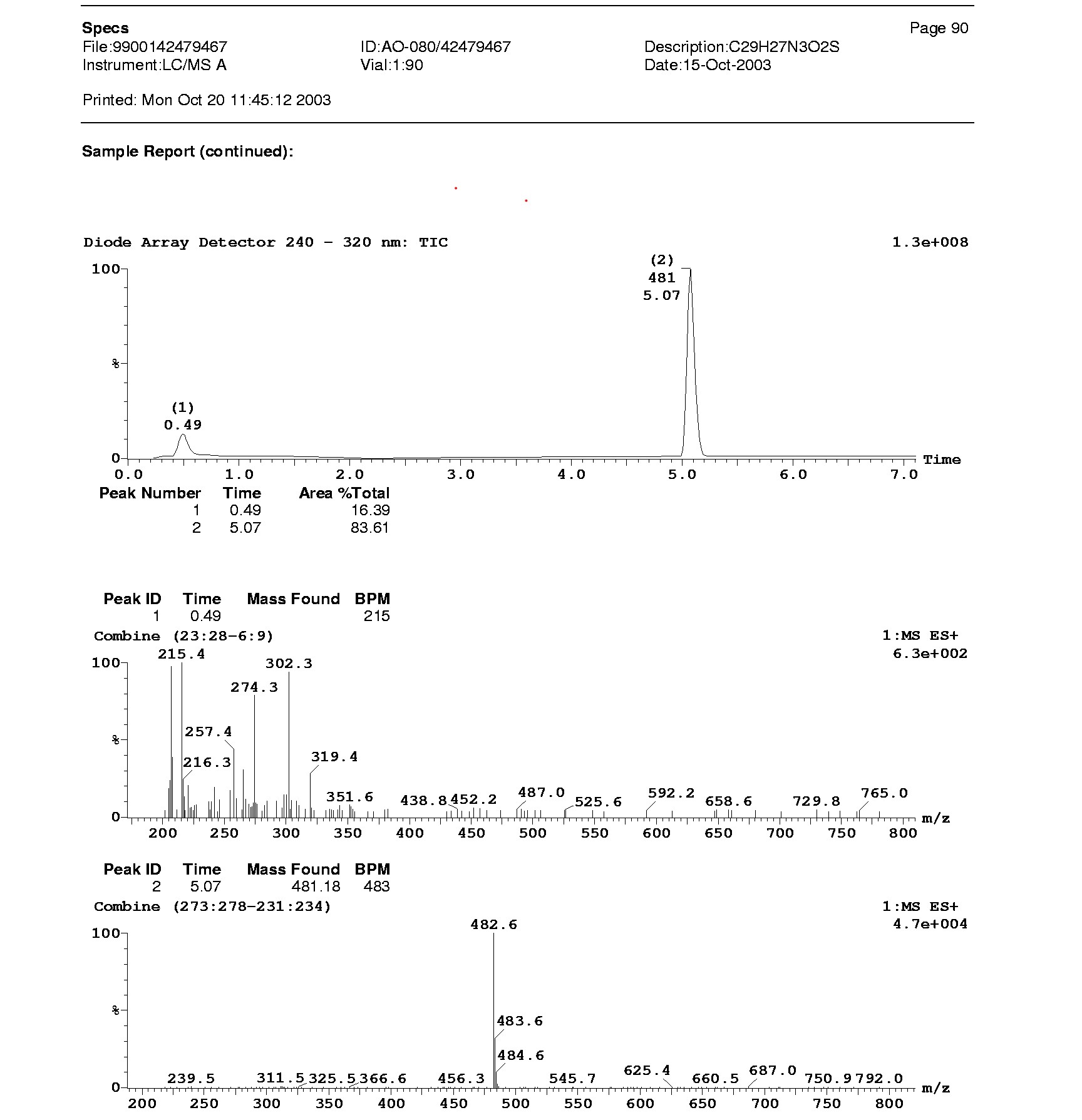

^1^H-NMR of AK-968/40390012

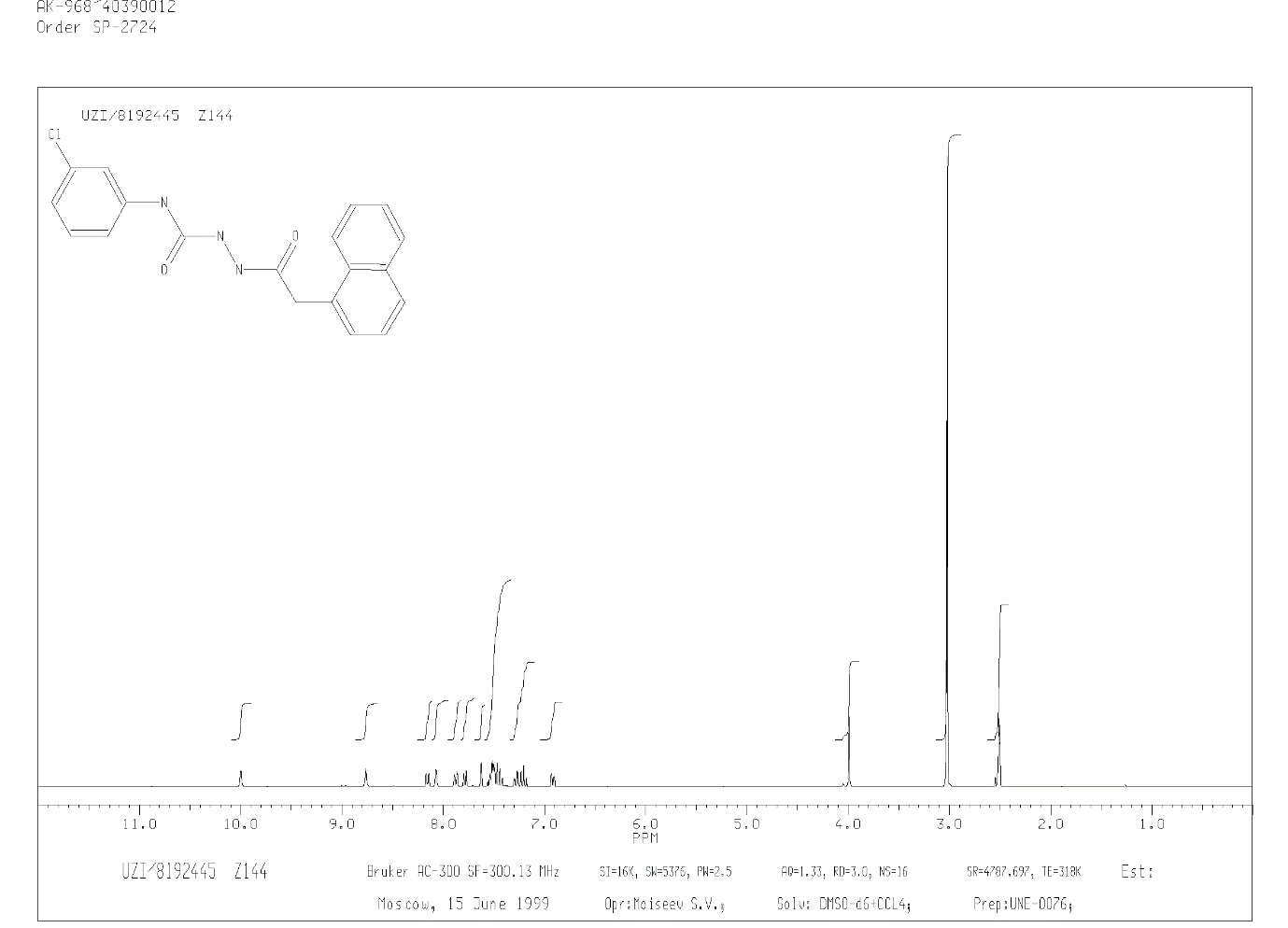

MS of AK-968/40390012

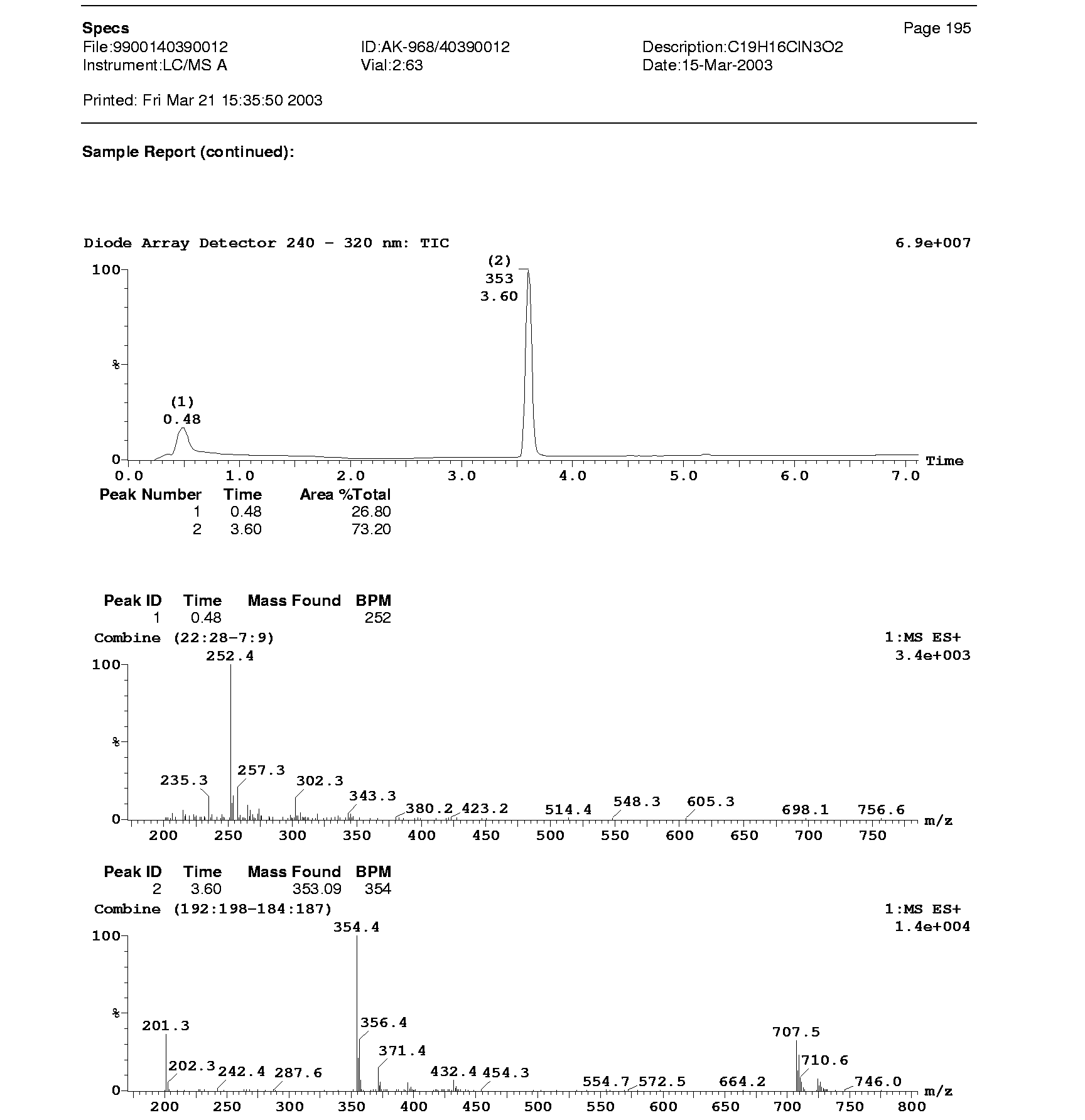

^1^H-NMR of Z384684494

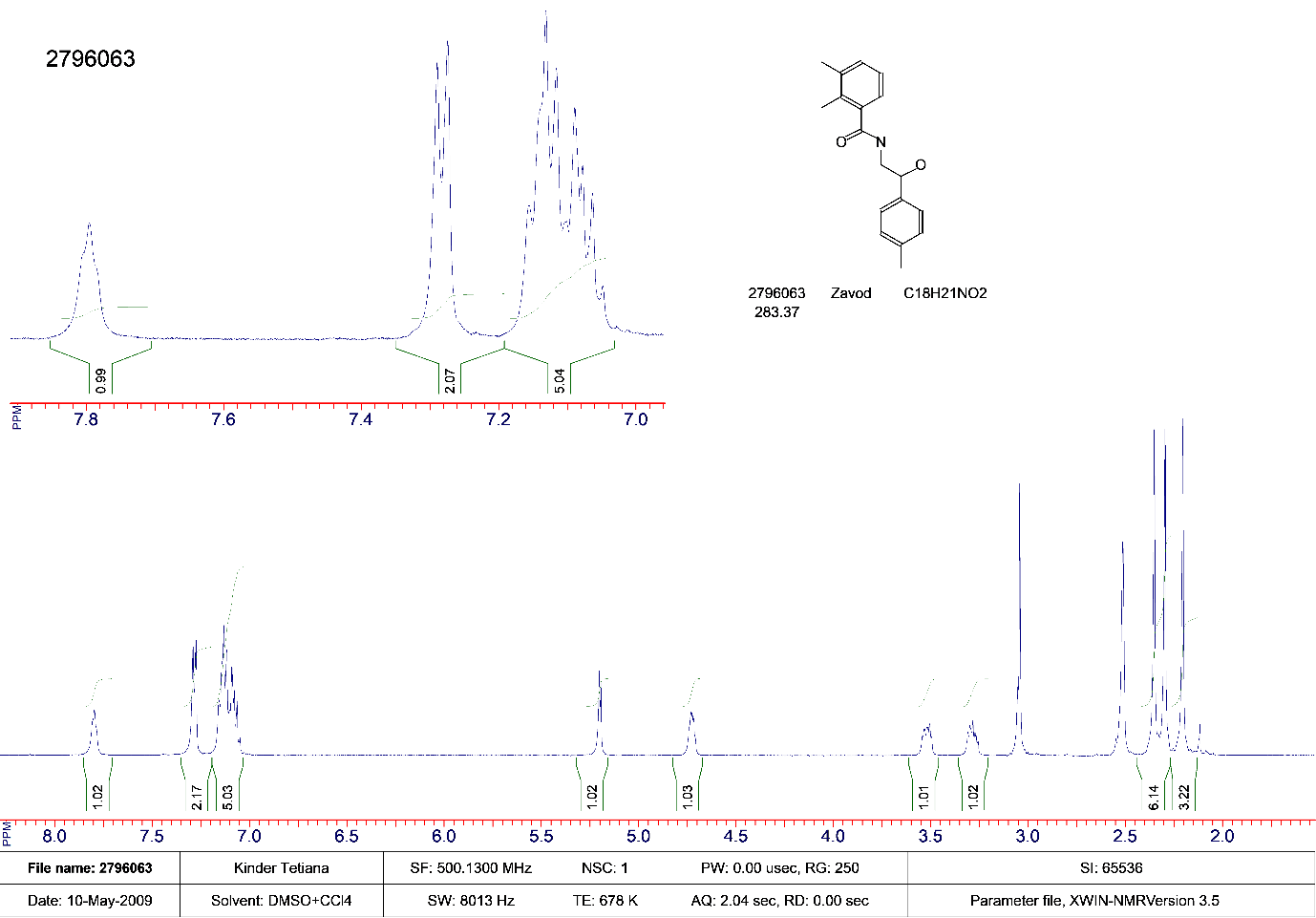

^1^H-NMR of AQ-149/43243694

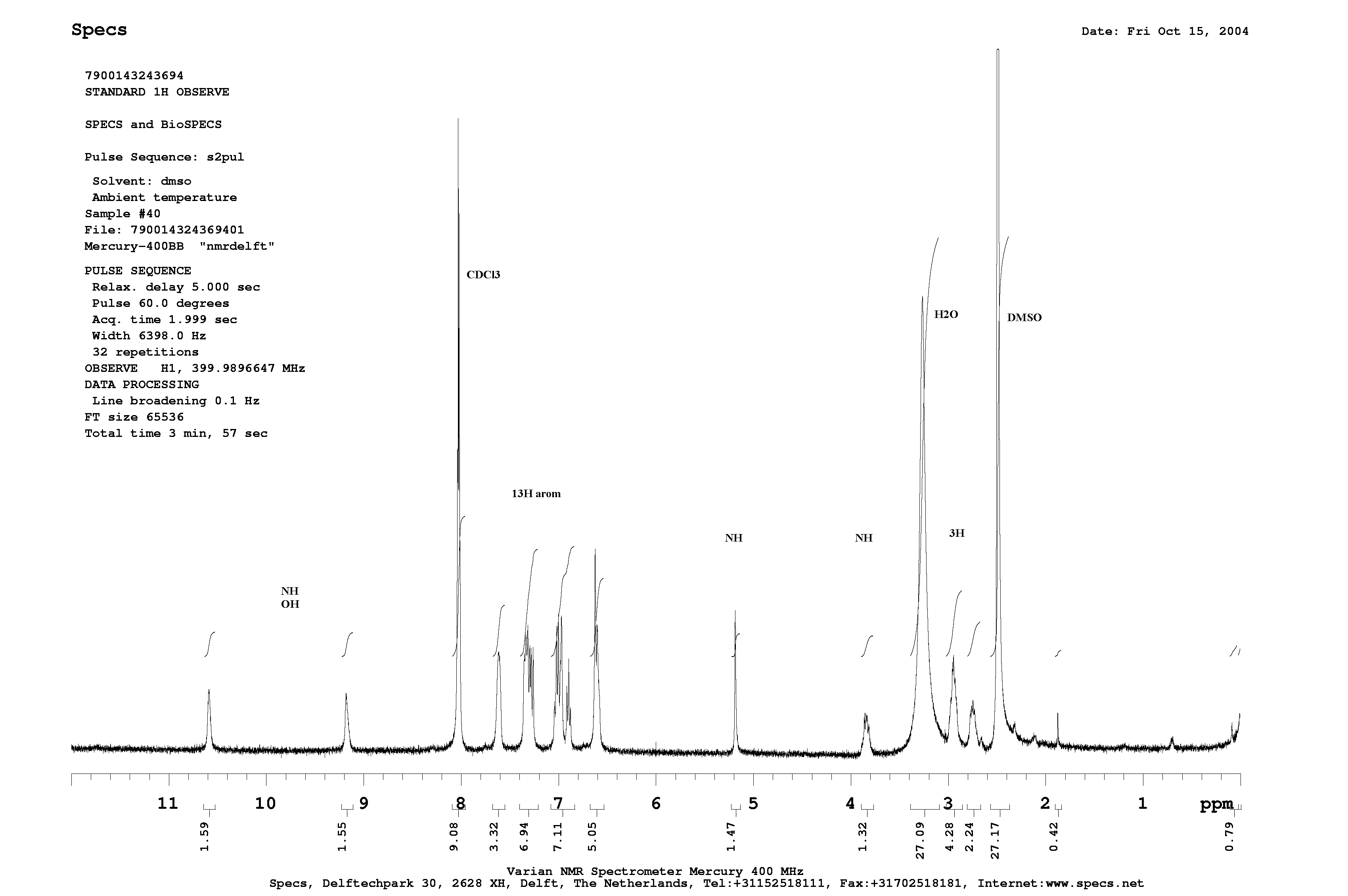

MS of AQ-149/43243694

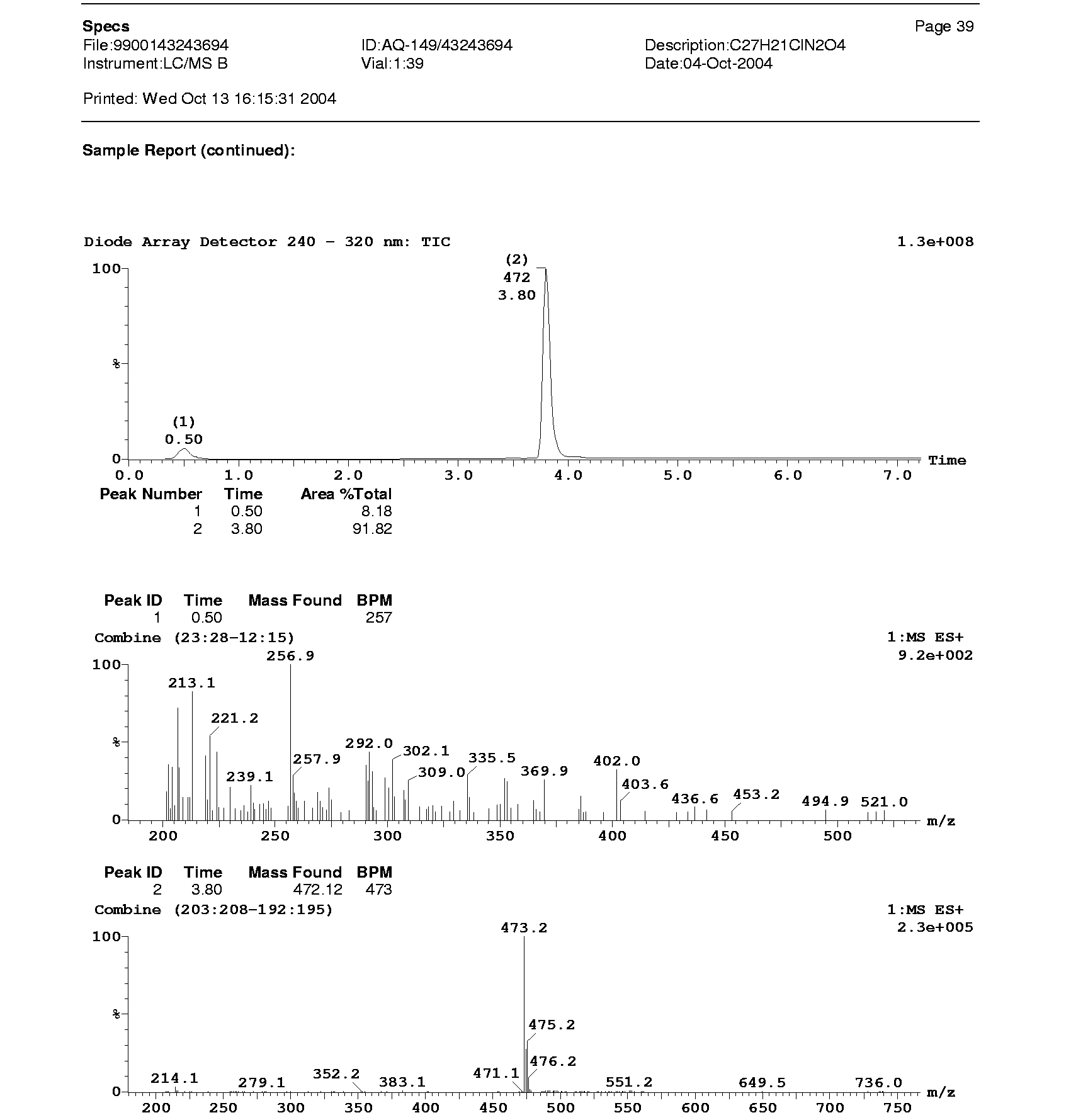

^1^H-NMR of E642-0121

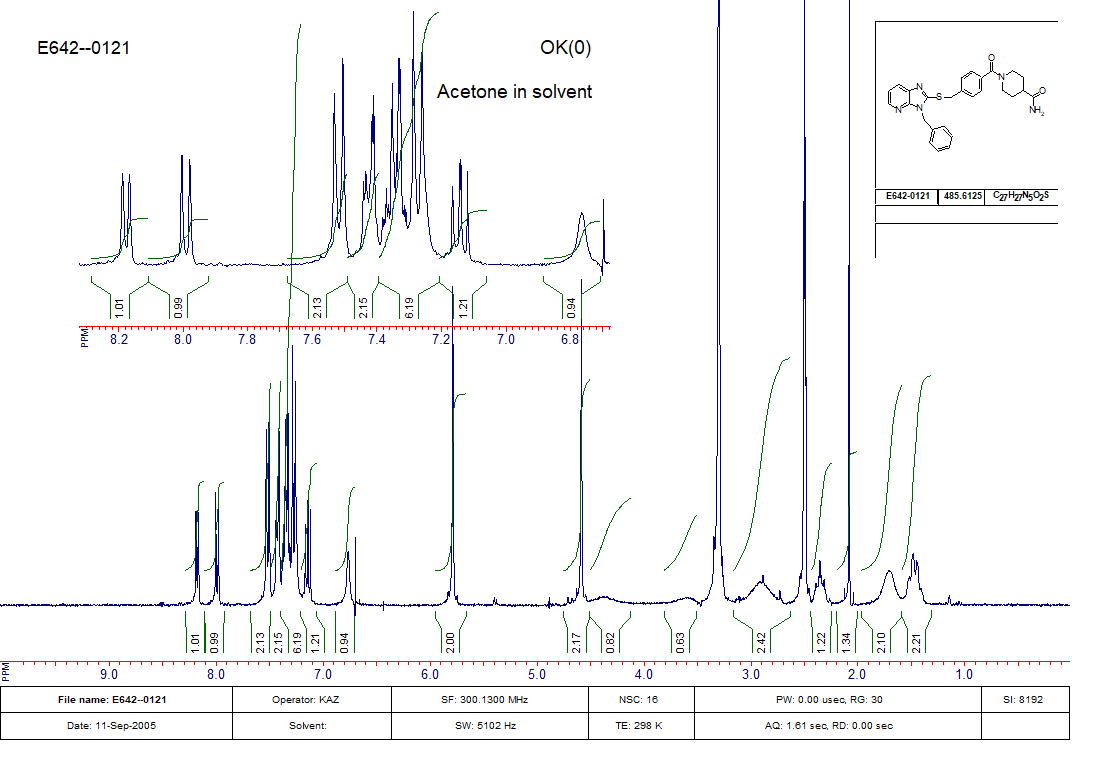

^1^H-NMR of AO-080/43441933

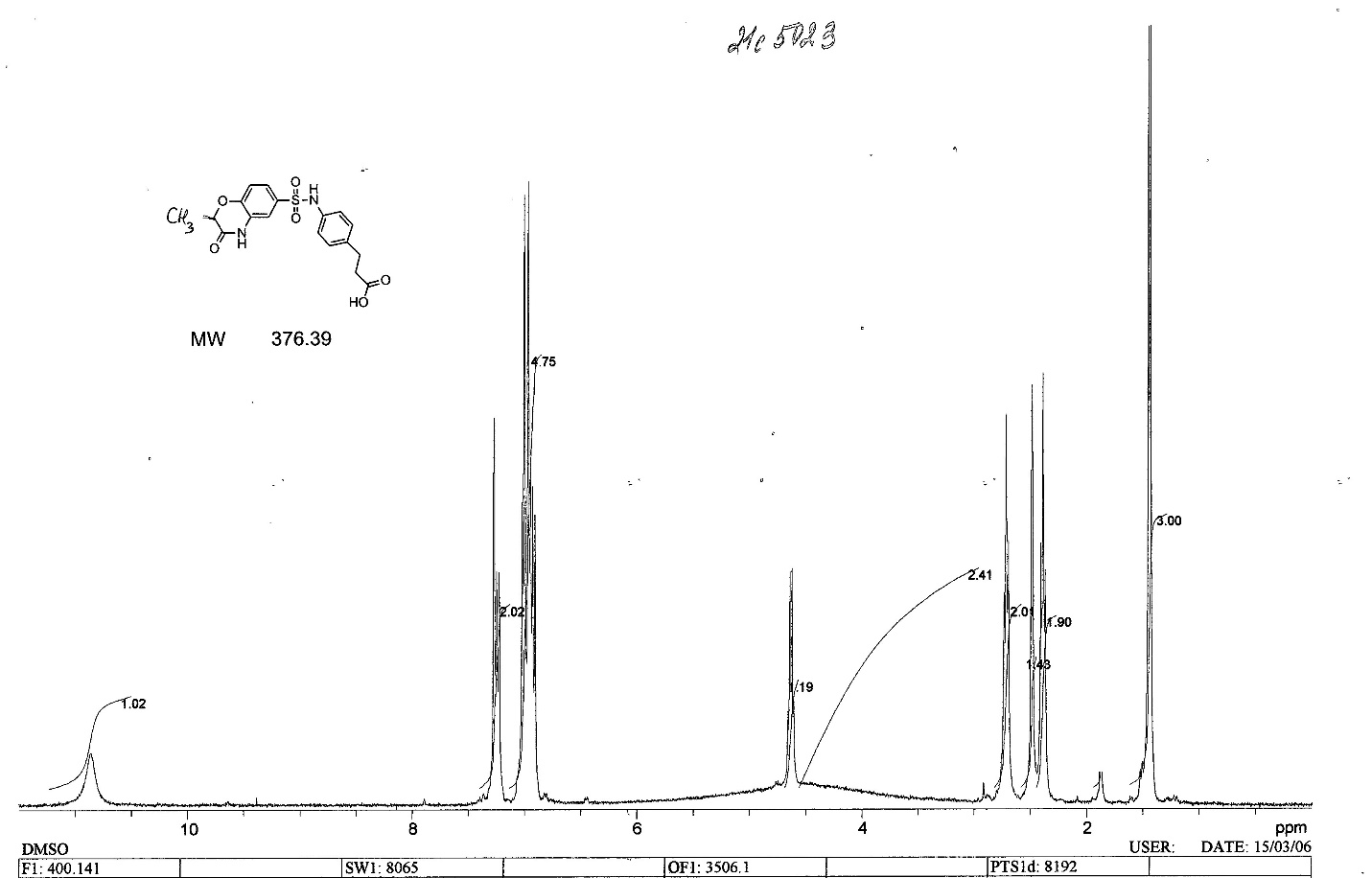

MS of AO-080/43441933

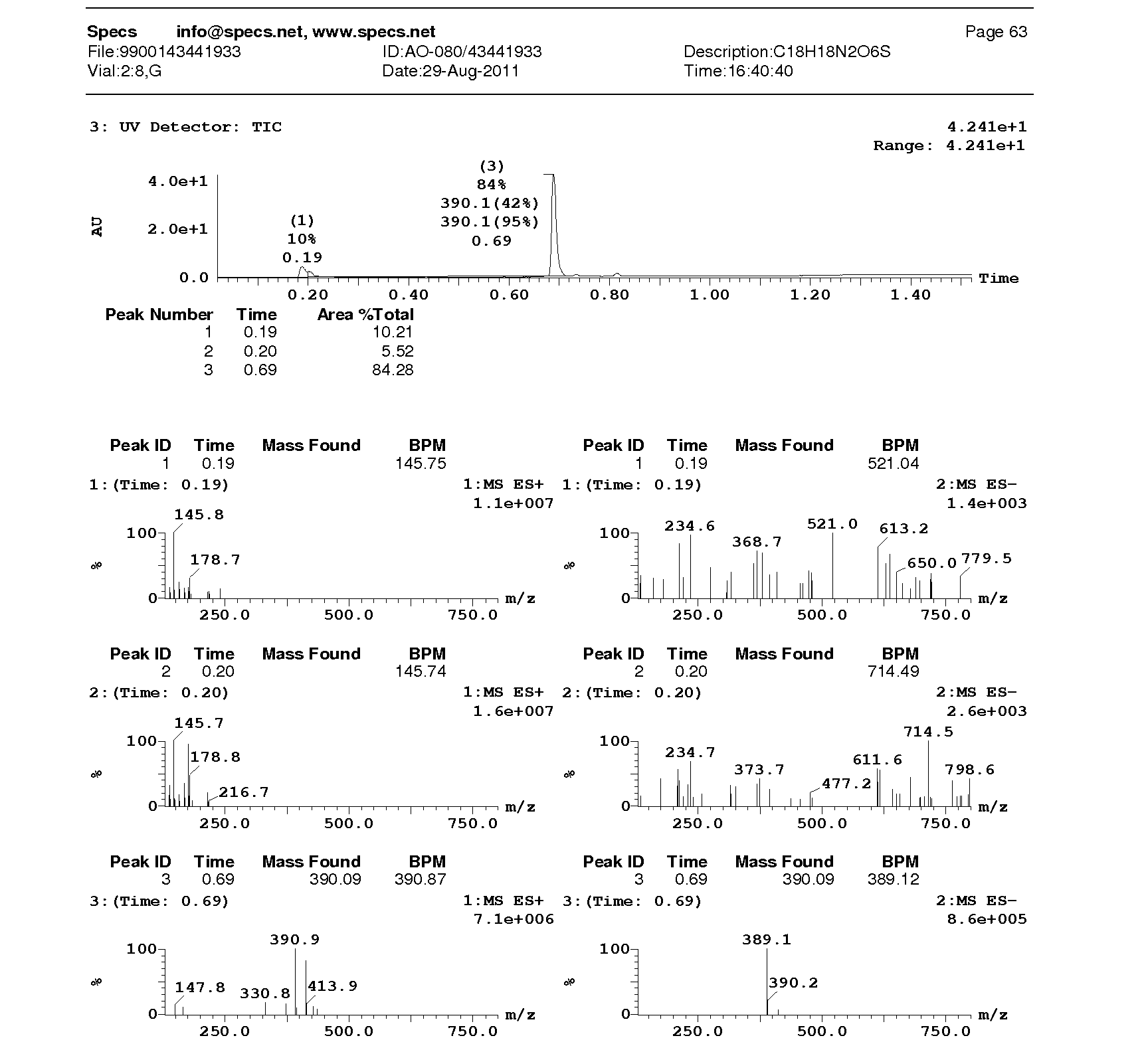

^1^H-NMR of AN-329/42613617

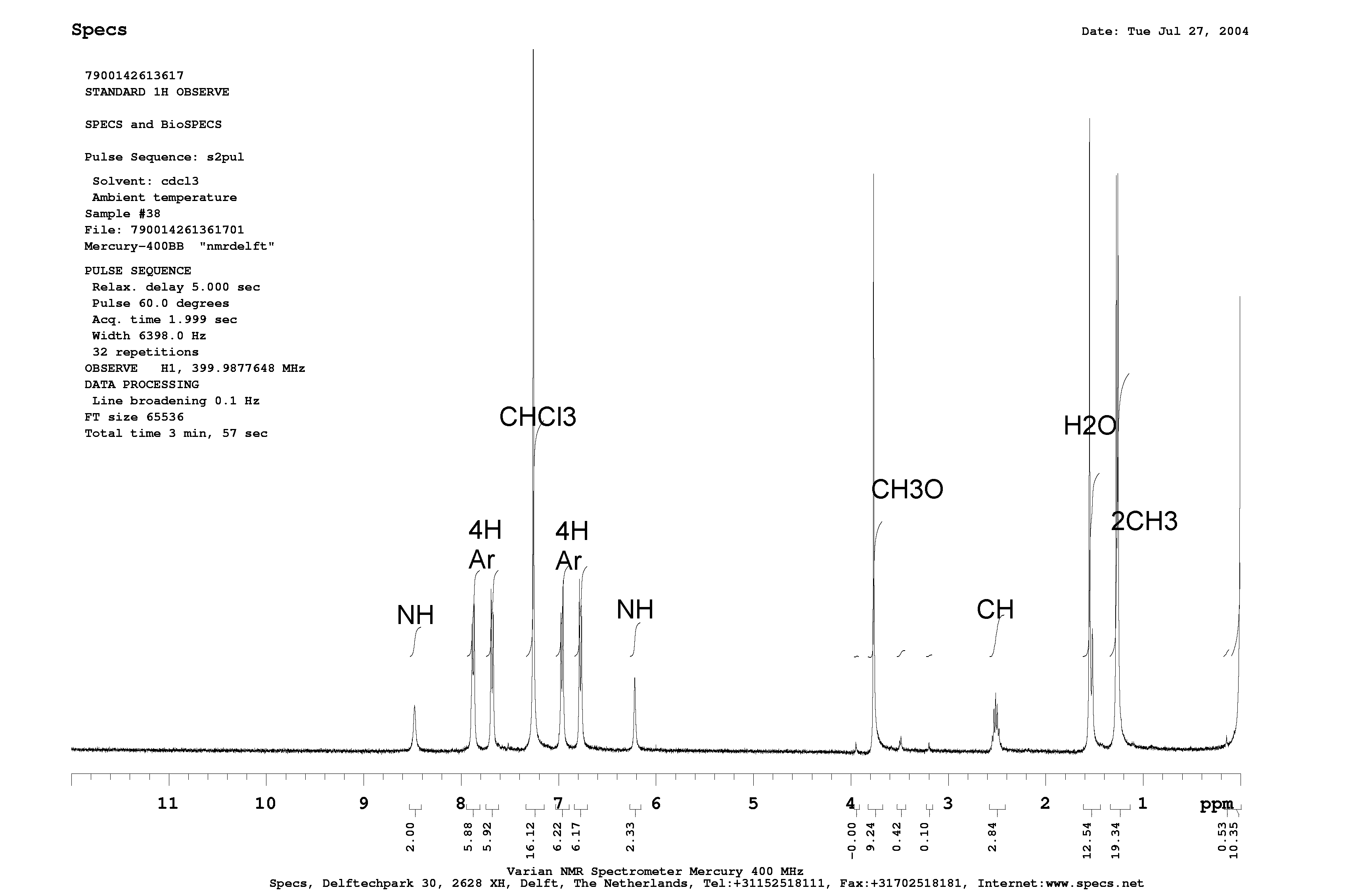

MS of AN-329/42613617

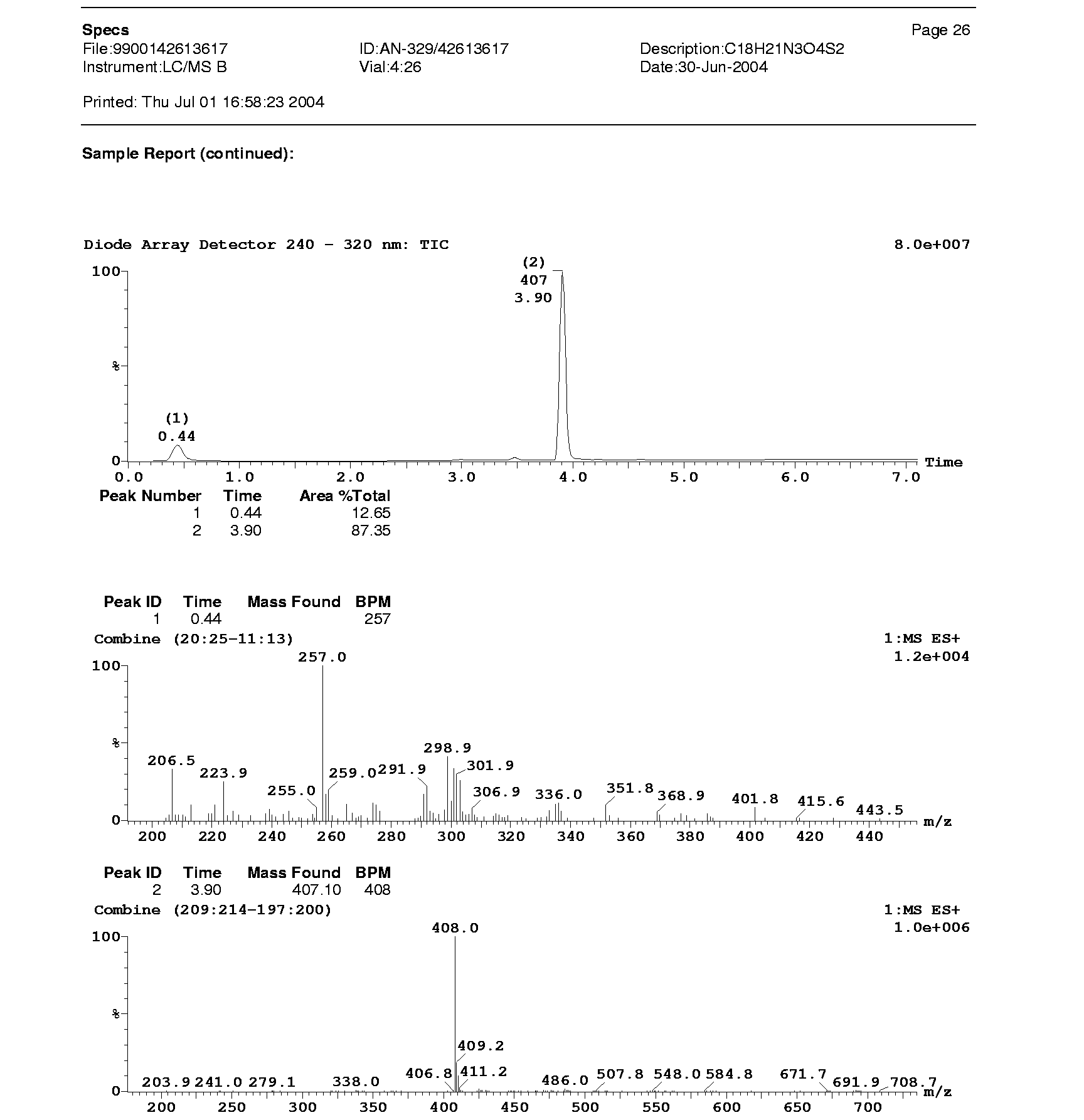

MS of AK-778/43420895

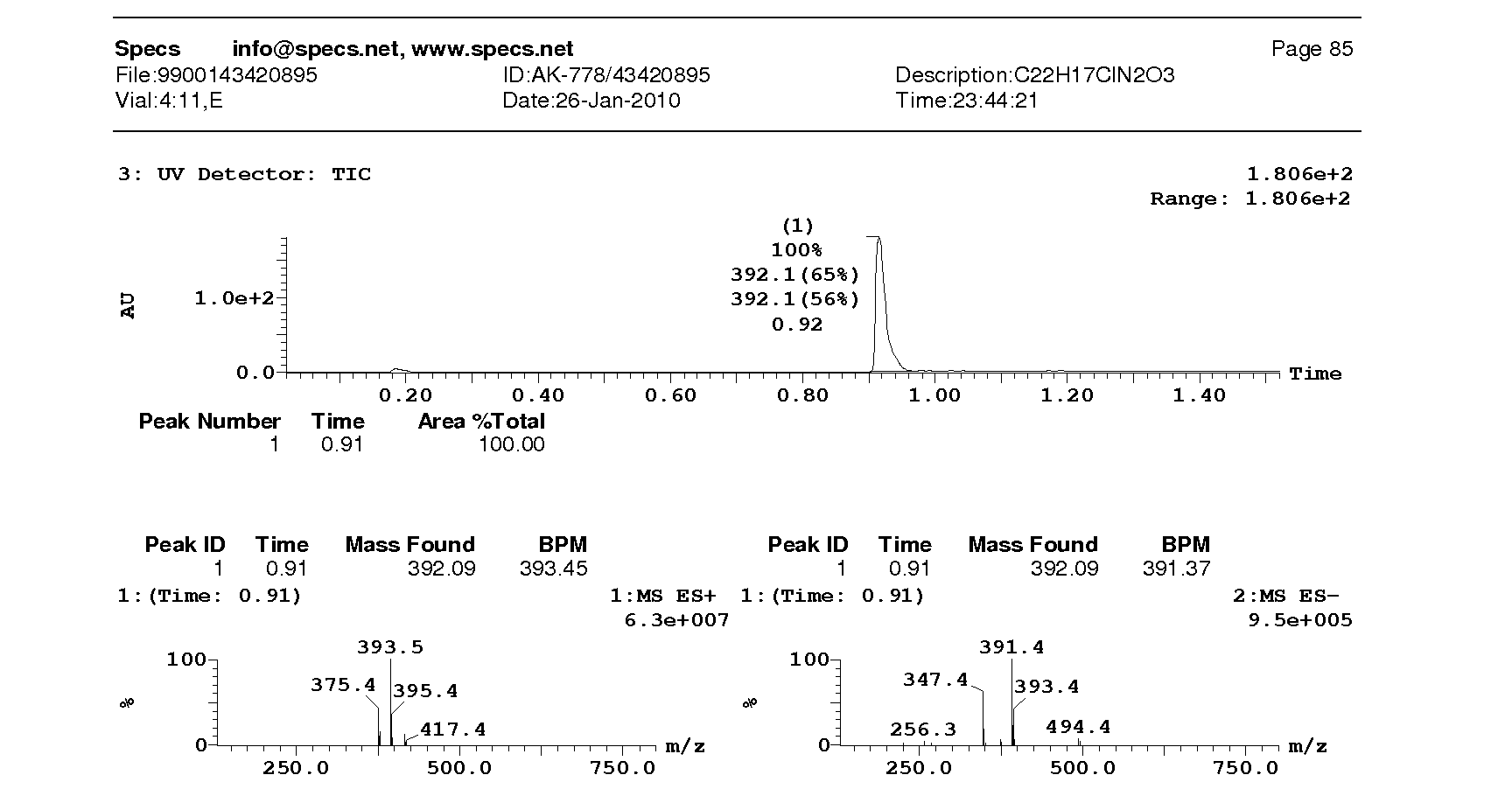

^1^H-NMR of F711-0605

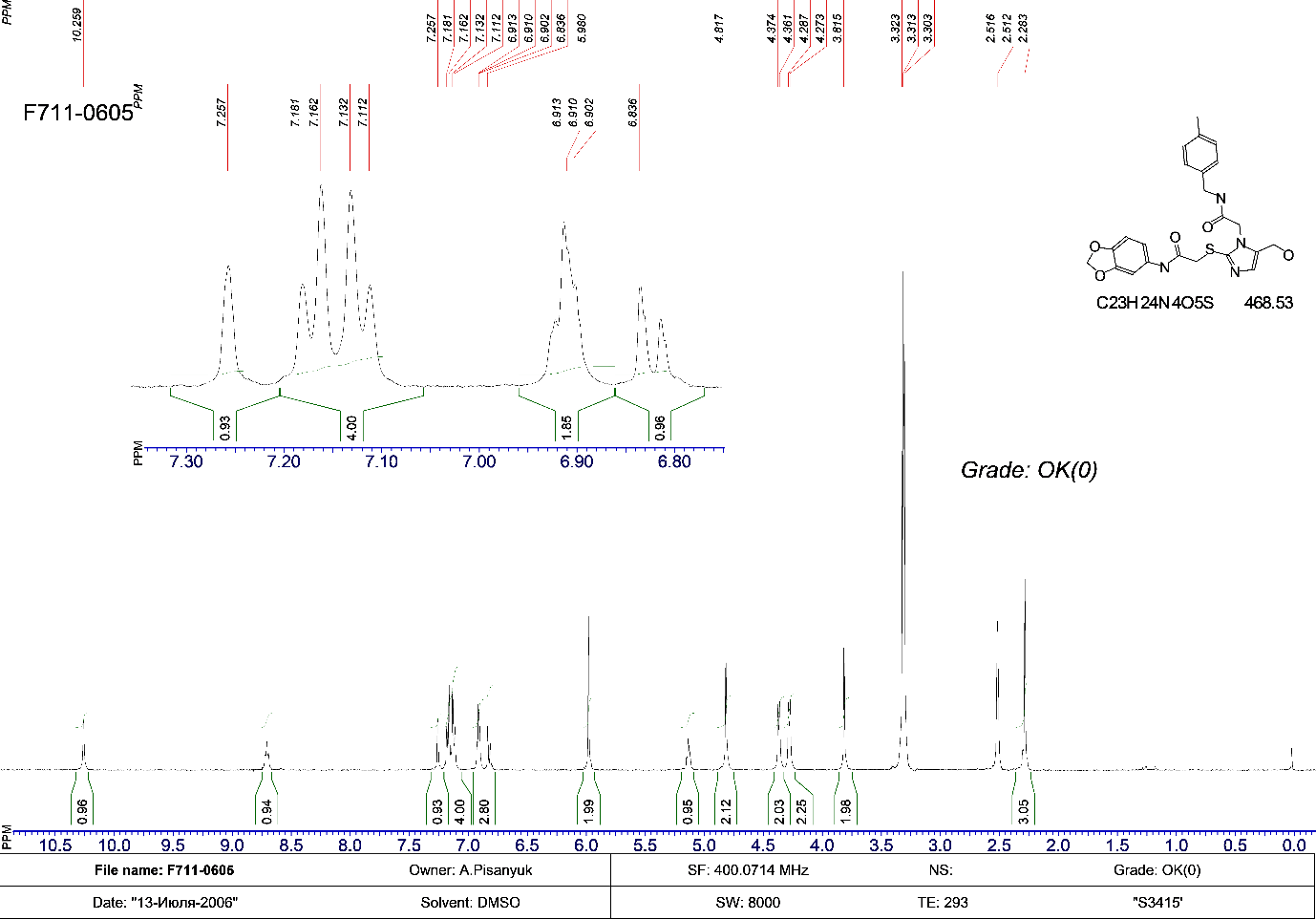

Table S1: SPR results between ID2 and AK-778-XXMU

| **Method** | **Ligand** | **Immobilized Lever （RU）** | **Analyte** | **Analyte Conc.**  **(μM)** | **ka (1/Ms)** | **kd (1/s)** | **KD (M)** | **Rmax (RU)** | **Chi² (RU²)** | **Kinetics model** |
| --- | --- | --- | --- | --- | --- | --- | --- | --- | --- | --- |
| CM5 | ID2 | 7943.3 | AK-778-XXMU | 100, 50, 25, 12.5, 6.25, 3.125 | 2.48E+01 | 3.19E-06 | 1.29E-07 | 83.6 | 0.23 | 1:1 binding |

Fig. S1: The response curve from SPR

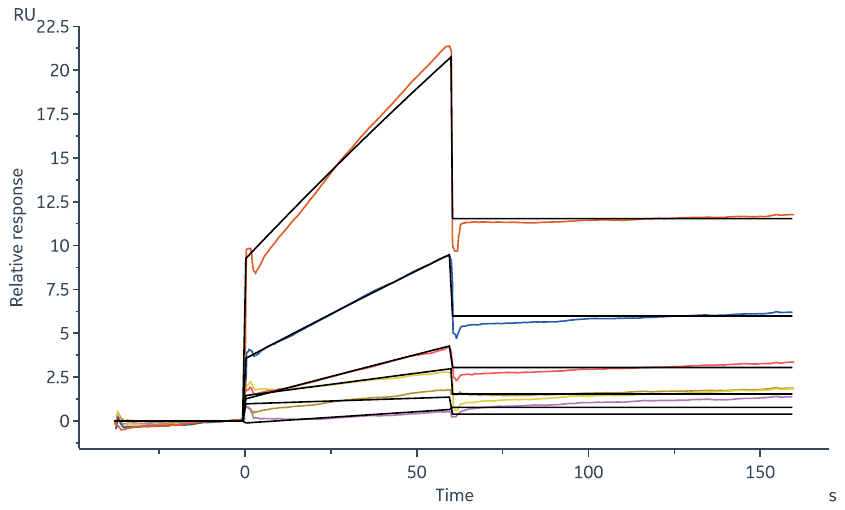
